# A family of lanthipeptides with anti-phage function

**DOI:** 10.1101/2024.06.26.600839

**Authors:** Helena Shomar, Florian Tesson, Marie Guillaume, Véronique Ongenae, Margot Le Bot, Héloïse Georjon, Ernest Mordret, Le Zhang, Gilles P. van Wezel, Daniel Rozen, Ariane Briegel, Séverine Zirah, Dennis Claessen, Yanyan Li, Aude Bernheim

**Affiliations:** Institut Pasteur, Université Paris-Cité, CNRS UMR 3525, Molecular Diversity of Microbes, Paris, France; INSERM, SEED, UMR 1284, Université Paris-Cité, Paris, France; INSERM, IAME, UMR 1137, Université Paris-Cité, Paris, France; Institute of Biology, Leiden University, Sylviusweg 72, 2333 Leiden, The Netherlands; Laboratory Molecules of Communication and Adaptation of Microorganisms (UMR 7245 CNRS-MNHN), Muséum National d’Histoire Naturelle, CP 54, 57 rue Cuvier, 75005, Paris, France; Generare Bioscience, Paris, France

## Abstract

Bacteria synthesize natural products to adapt to their environment, where phage-bacteria interactions play a crucial role in bacterial ecology. Although a few natural products have been shown to protect bacteria from phage infection, the prevalence and diversity of chemical anti-phage defense remain largely unexplored. Here, we uncover a novel family of over 2000 lanthipeptide biosynthetic gene clusters from Actinobacteria that participate in anti-phage defense, which we named lanthiphages. Lanthiphages colocalize with other anti-phage systems in defense islands. We demonstrate that native lanthiphage expression protects the model strain *Streptomyces coelicolor* against diverse phages. Heterologous expression of four additional lanthiphage pathways shows that the anti-phage function is conserved across this family of biosynthetic gene clusters. Finally, we demonstrate that lanthiphage expression leads to the production of a novel compound and alters phage transcription. Our findings highlight that biosynthetic gene clusters with anti-phage functions can be successfully identified through genomic analysis. This work paves the way for the systematic mining of anti-phage natural products, which could constitute a novel reservoir of antiviral drugs.

## Introduction

Bacteria produce chemical compounds, known as natural products, to adapt to their abiotic and biotic environments^1,2^. While thousands of natural products, such as antibiotics and anticancer agents, have been discovered and characterized in relation with their clinical^3,4^ and economic value^5,6^, the biological function of most natural products remains unknown. Phage-bacteria interactions play a key role in bacterial ecology^7,8^. Reports from the 1940s noted that *Streptomyces* culture extracts can inhibit phage replication^9,10^, but it was only recently demonstrated that a few known natural products, such as DNA intercalating agents^11^ (daunorubicin, doxorubicin), aminoglycosides^12^ (apramycin, kanamycin), and modified nucleotides^13^ protect bacteria from phages. However, the prevalence and diversity of anti-phage “chemical defense” remains largely unexplored.

Recent advances in genomics have enabled the prediction of genes potentially involved in natural product synthesis^14–16^ or anti-phage defense^17–19^. Here we set out to integrate these predictions to identify new anti-phage chemical defense mechanisms. The majority of microbial natural products are synthesized by metabolic pathways encoded in organized groups of operons called biosynthetic gene clusters (BGCs)^14^. As only 3% of predicted BGCs specify a characterized compound^20^, we reasoned that uncharacterized BGCs could offer a valuable reservoir of anti-phage compounds. Anti-phage systems cluster in genomic regions called defense islands^21^. Leveraging the colocalization of known anti-phage genes has allowed the discovery of dozens of novel anti-phage systems^7^. Here, we hypothesized that analyzing genomic colocalization of BGCs and defense islands could uncover BGCs with anti-phage function.

Lanthipeptides are the largest and most diverse class of ribosomally synthesized and post-translationally modified peptides (RiPPs), a superfamily of natural products^22,23^. Lanthipeptides are synthesized from an encoded precursor peptide that undergoes posttranslational modifications catalyzed by a set of specialized enzymes, resulting in the formation of lanthionine rings^23^ (Fig. 1a). While the number of discovered lanthipeptides has increased in recent years, little is known about the diversity of their biological functions and whether they might be involved in anti-phage defense^24^. Here, we describe a novel family of lanthipeptide BGCs from Actinobacteria that colocalize with known anti-phage defense systems and show they confer anti-phage activity.

**Figure 1.**
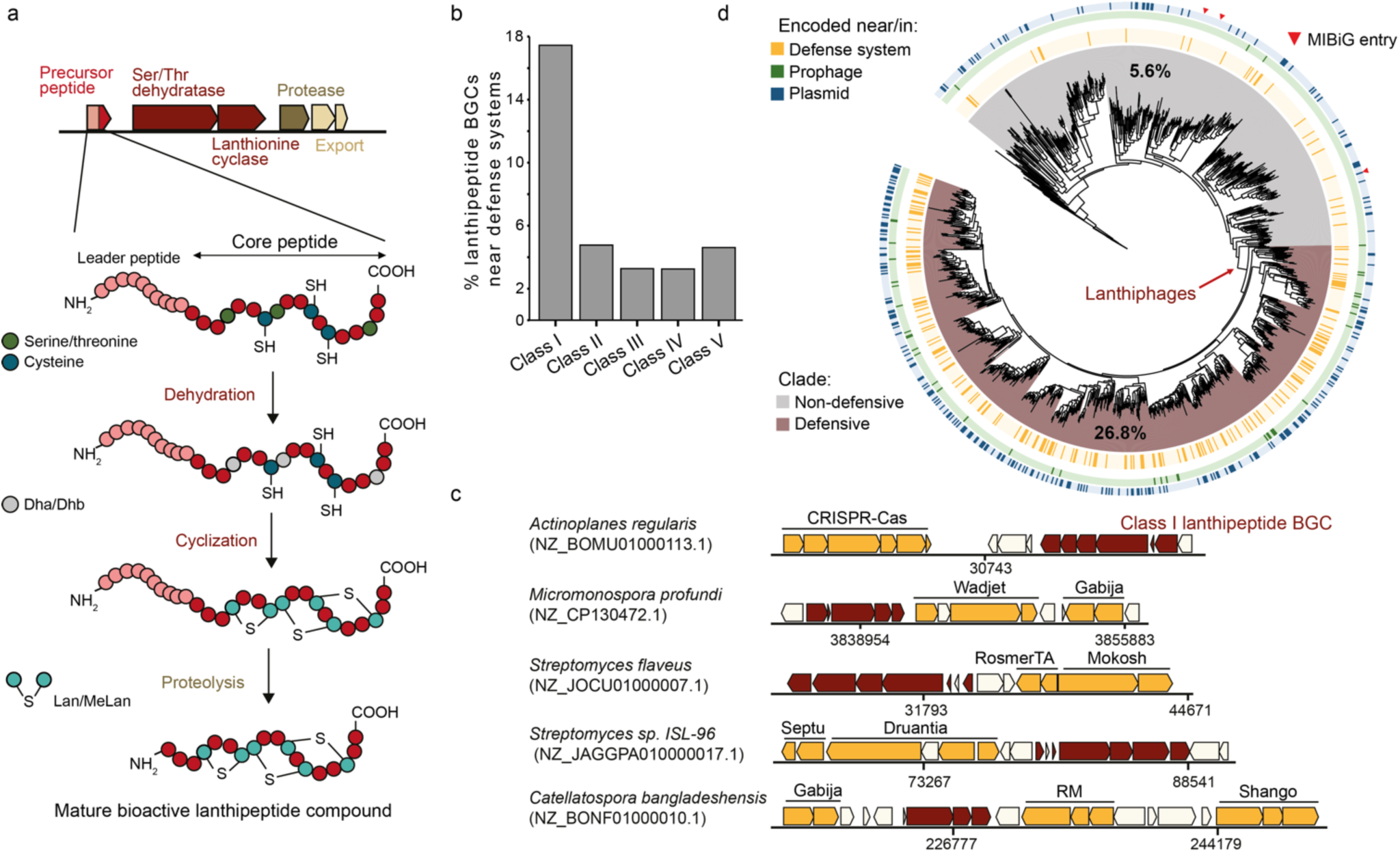
Detection of a clade of class I lanthipeptide BGCs encoded in defense islands and mobile genetic elements. **a.** Schematic representation of lanthipeptide biosynthesis^28^. Lanthipeptides are produced from a genetically encoded precursor peptide, which consists of a N-terminal leader and a C-terminal core peptide. The core biosynthetic enzymes post-translationally modify the core peptide by catalyzing the dehydration of serine and threonine residues, yielding respectively dehydroalanine (Dha) and dehydrobutyrine (Dhb), followed by addition of cysteine thiols onto these residues to form lanthionine (Lan) or methyllanthionine (MeLan) rings. Proteolytic cleavage of the leader peptide yields the mature bioactive lanthipeptide compound. Specific protease and/or ABC transporters are often encoded in lanthipeptide BGCs, and sometimes additional enzymes that catalyze secondary modifications. **b.** Differential enrichment of different lanthipeptide BGC classes located near defense systems in actinobacterial genomes. **c.** Representative instances of lanthiphages (in red) and their genomic context. Genes encoding known defense systems are shown in yellow. RM: restriction modification. **d.** Phylogenetic tree of actinobacterial class I lanthionine cyclases, rooted using lanthionine cyclases from class II lanthipeptide BGCs as an outgroup. Colored rings describe the genetic context of BGCs. For each clade, the percentage of BGCs that are located near known defense systems is presented. Pink triangles indicate characterized BGCs from the MIBiG database that synthesize nocathioamide A (BGC0002120), planosporicin (BGC0000544) and microbisporicin A2 (BGC0000529).

## Results

### A family of class I lanthipeptide BGCs is enriched in defense islands

To mine lanthipeptide BGCs located in defense islands, we analyzed 29,401 genomes from the BGC-rich phylum Actinobacteria using antiSMASH^25^ and identified 13,701 unique BGCs predicted to be involved in lanthipeptide production. AntiSMASH employs a rule-based approach to identify various types of BGCs, and provides detailed analyses for different lanthipeptide BGC classes, including predictions of core biosynthetic enzymes, accessory enzymes, and encoded core peptides (Fig 1a). Next, we used DefenseFinder^26^ to detect known anti-phage defense systems within the vicinity of each BGC (+/- 23 genes, see Methods). We found 1,200 lanthipeptide BGCs located near at least one defense system (8.8% of the total). Class I lanthipeptide BGCs were more frequently found near defense systems (17.7%, overall next to 83 different defense system subtypes) compared to other classes of lanthipeptide BGCs (between 4%-5%, Fig 1b-c) similar to what is expected for a random gene (4.7%).

To further characterize the diversity of class I lanthipeptide BGCs located near anti-phage systems, we built a phylogenetic tree using 1,031 representative lanthionine cyclases from class I lanthipeptide BGCs and mapped the presence of known anti-phage systems in their vicinity (Fig 1d, Supplementary Table 1). This analysis revealed a specific clade of BGCs enriched in defense islands (defensive clade, light pink) with 26.6% BGCs located near known anti-phage systems, compared to 5.6% in other class I lanthipeptide BGCs (non-defensive clade, gray) (Fig 1d). BGCs from the defensive clade are also more frequently found on prophages or plasmids (Fig 1d, Supplementary Fig 1a). All characterized lanthipeptide BGCs from the MIBiG reference database^27^ (n=64) are located outside the defensive clade (Fig 1d, Supplementary Fig 1b). No additional defense-associated lanthipeptide clades were identified in other lanthipeptide classes (Supplementary Fig 1b). These findings suggest the unique emergence of a clade of class I lanthipeptide BGCs enriched in defense islands. Given that BGCs from the defensive clade are uncharacterized and associated with anti-phage defense systems, we hypothesized that they could have an anti-phage function and we therefore named them “lanthiphages”.

### Expression of a native lanthiphage protects against phage infection

Genomic analysis unveiled the presence of a lanthiphage in the genome of the model strain *Streptomyces coelicolor* M145 (Fig 2a). This lanthiphage (lanthiphage_Sco) is one of the two remaining cryptic and uncharacterized BGCs among the 27 BGCs found in this model organism^29^. It contains genes encoding for the typical lanthipeptide core biosynthetic enzymes, two precursor peptide genes with identical predicted core peptides and two genes for accessory proteins that have been previously described^30,31^(Fig 2a).

**Figure 2.**
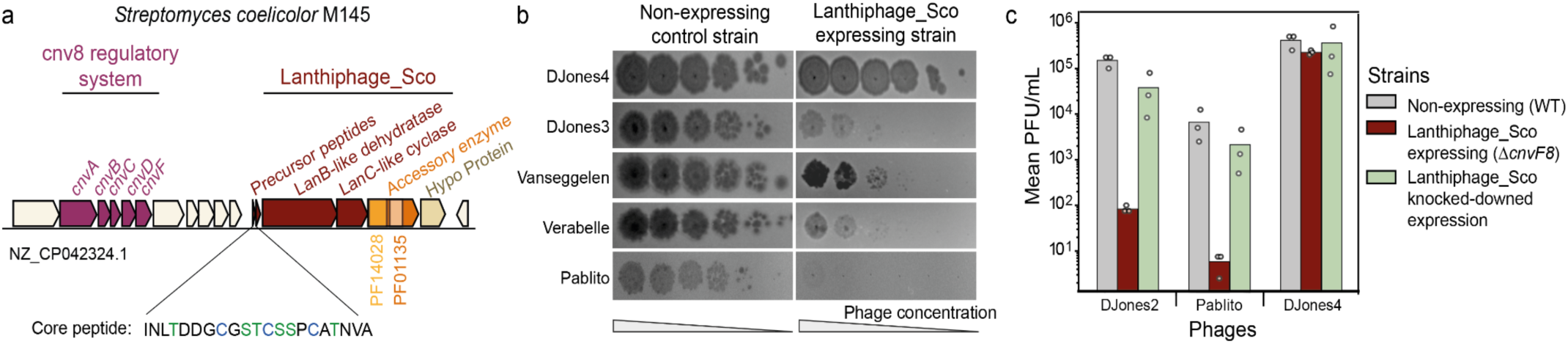
Expression of a native lanthiphage confers anti-phage resistance in *S. coelicolor*. **a.** Schematic representation of the genomic region of *S. coelicolor* M145 that contains lanthiphage_Sco and the nearby conservon regulatory system Cnv8. Depicted in dark red are the predicted precursor peptide genes and genes for core biosynthetic enzymes, namely LanB-like serine/threonine dehydratase and LanC-like lanthionine cyclase. In addition, lanthiphage_Sco encodes two accessory proteins, one enzyme with L-isoaspartate O-methyltransferase (PF01135) and thiopeptide-type bacteriocin biosynthesis (PF14028) domains, and a hypothetical protein (structurally similar to tetratricopeptide-repeat proteins). Highlighted in color are residues of the core peptide that are post-translationally modified and participate in lanthionine ring formation: serine/threonine in green, cysteine in blue. **b.** Plaque assay results shown for five phages from the DJones collection on the lanthiphage-expressing strain (*ΔcvnF8*) compared to the non-expressing WT *S. coelicolor* strain. Images are representative of three biological replicates. Quantifications for all phages are found in Supplementary Fig 3. **c.** Effect of knocking-down lanthiphage_Sco genes on the anti-phage resistance of the lanthiphage_Sco expressing strain. Using the Cumate-Based Inducible CRISPRi (CUBIC) system^32^, we designed guide RNA sequences (gRNA) to knock down the expression of the LanB-like enzyme (strain CUBIC_DH, green bars). Bars represent an average of three biological replicates, with individual data points overlaid. Similar results are observed using gRNA sequences that target both precursor peptide genes simultaneously (Supplementary Fig 3b).

To test the hypothesis that lanthiphage_Sco has an anti-phage function, we investigated whether its native expression would confer phage resistance. A recent study demonstrated that a regulatory system known as conservon 8 (Cvn8) regulates the expression of lanthiphage_Sco located 4.3 kb downstream^33^ (Fig2a). Single knock-out mutants of the *cnv8* operon displayed acute upregulation of the lanthiphage_Sco genes, indicating that the Cvn8 system strongly represses lanthiphage_Sco expression^33^. Therefore, we compared the susceptibility of lanthiphage_Sco expressing (Δ*cvnF8* knock-out mutant) and non-expressing (wild-type) strains against a set of diverse phages that infect *S. coelicolor* M145. For this purpose, we built a collection of 11 phages that infect *Streptomyces* species, isolated from environmental soil samples named the Djones collection (after Doris Jones, see Supplementary Text 1, see Methods, Supplementary Fig 2). Plaque assay results show a significant reduction in plaque formation in the lanthiphage_Sco expressing strain compared to the non-expressing train for 10 of 11 tested phages. (Fig2b, Supplementary Fig 3a). For instance, we observed a 5,000-fold reduction of phage DJones3 titer between lanthiphage-expressing and non-expressing strains (Fig2b, Supplementary Fig 3a). To further validate the anti-phage function of lanthiphage_Sco, we knocked down the expression of lanthiphage_Sco genes in the expressing strain (either both precursor peptide genes simultaneously or the LanB-like enzyme in *ΔcvnF8*, see Methods). In all cases, we observed the disappearance of phage defense (Fig2c, Supplementary Fig 3a), indicating that lanthiphage_Sco expression is essential for anti-phage activity.

### Genomic characterization of lanthiphages

To assess the prevalence of lanthiphages beyond Actinobacteria, we detected their presence in 22,793 prokaryotic genomes, including 20,540 non-actinobacterial genomes. Lanthiphages are found exclusively in Actinobacteria but distributed across diverse genera (Supplementary Fig 4). Among the analyzed genomes, 29% of *Streptomyces,* 57% of *Salinispora* and 83% of *Frankia* species contain lanthiphages. The number of lanthiphages per genome (Supplementary Fig 4) varies from zero to six (for instance in strain *Frankia casuarinae CcI3)*.

To investigate the diversity of lanthiphages, we built a phylogeny of all lanthiphage-associated lanthionine cyclases (n=2,064) (Fig 3a). Lanthiphages group in ten major clades (Fig 3a) and display diverse genomic architectures (Fig 3b, Supplementary Fig 5a, Supplementary Table 1). We integrated both these features (phylogeny and genomic architecture) to define seven lanthiphage types (Fig 3b, Supplementary Fig 5a, Supplementary Table 1). All lanthiphages, including lanthiphage_Sco (Fig 2a), encode the conserved LanB-like and LanC-like core enzymes, here named LphB and LphC respectively, as well as additional accessory enzymes named LphD (PF14028 domain), LphE (PF01135 domain), LphDE (fused PF14028 and PF01135 domains) and LphF (hypothetical protein) (Fig 3b). Although the functions of LphD, LphE, LphDE, and LphF are unknown, their conservation across lanthiphages and rare presence in non-defensive class I lanthipeptide BGCs (Supplementary Fig 1c) suggest they play an important role in lanthiphage biochemistry. Lanthiphages are also more frequently associated with conservon systems (3.8%) than class I lanthipeptide BGCs from the non-defensive clade (0.7%) (Supplementary Fig 1d, Fig 3a).

**Figure 3.**
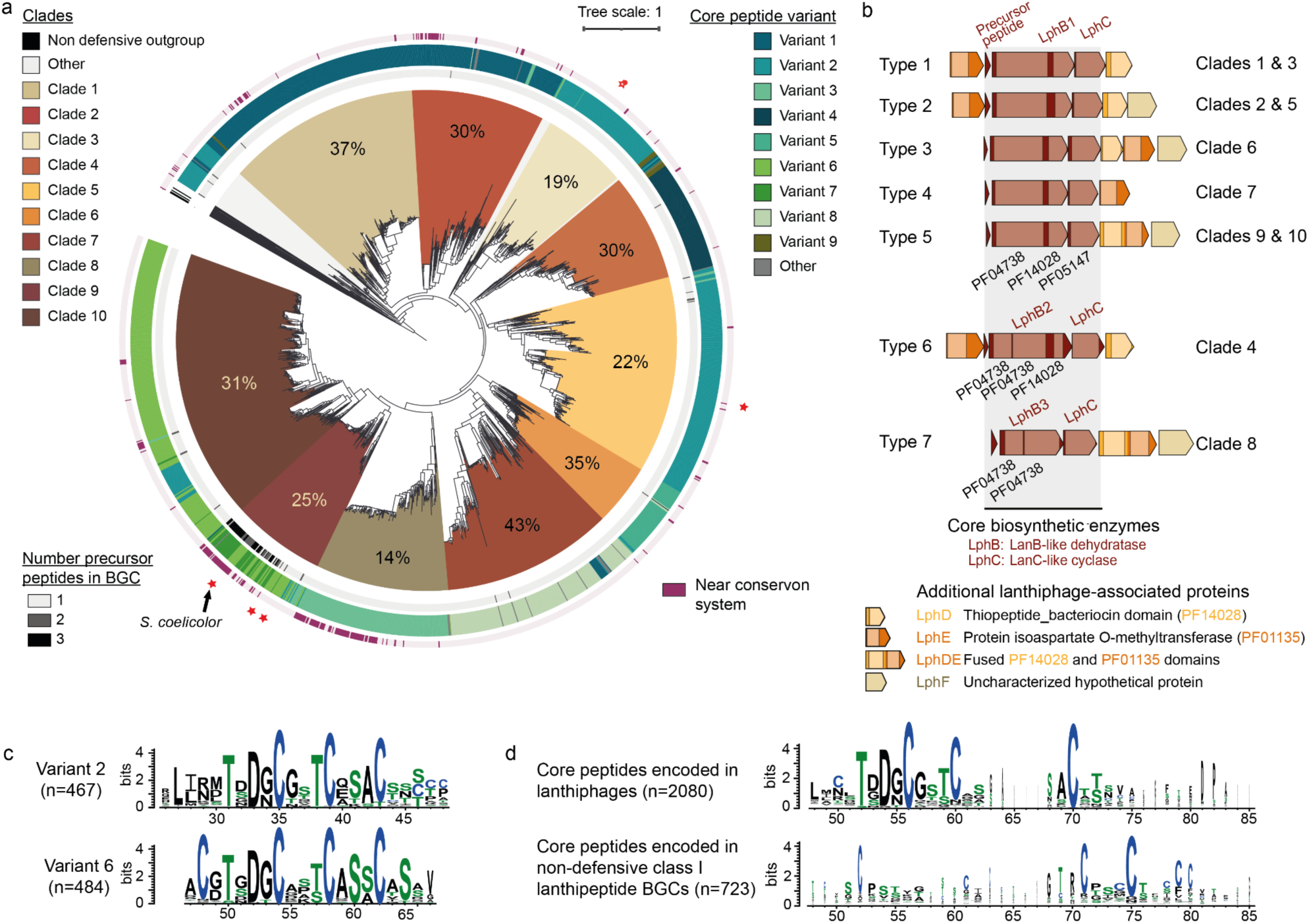
Genomic characterization of lanthiphages. **a.** Phylogenetic tree of class I lanthipeptide BGCs from the lanthiphage family using their respective lanthionine cyclase sequences (n=2064). The sequences of lanthionine cyclases from class I lanthipeptides from the previously defined non-defensive clade were used as an outgroup. For each clade, the percentage of BGCs that are located near known defense systems is presented. Red stars indicate BGCs that were tested experimentally with confirmed anti-phage activity in this study. **b.** Classification of lanthiphages based on their genomic architecture and associated enzymes. **c.** Sequence logo representation of multiple sequence alignment of predicted core peptide variants, illustrating the relative frequency of conserved amino acids. Highlighted in color are residues that are post-translationally modified and participate in lanthionine ring formation: serine/threonine in green, cysteine in blue. **d.** Sequence logo representation of multiple sequence alignment of predicted core peptides from class I lanthipeptide BGCs: from the lanthiphage family, and from the non-defensive clade.

To further explore the diversity of compounds synthesized by lanthiphages, we analyzed the amino acid sequence of their predicted core peptides, which define the molecular scaffold of the produced lanthipeptides (Fig 1a). We identified 844 unique conserved core peptides encoded in lanthiphages (2018 in total), classified in 9 variants based on sequence similarity (Fig 3a,c, Supplementary Fig 5b, Supplementary Table 2, see Methods). 67 lanthiphages (predominantly from clade 9) encode multiple precursor peptide genes with distinct core peptide variants. Conservation analysis of all lanthiphage-associated core peptides revealed conserved amino acid motifs absent in core peptides from non-defensive class I lanthipeptide BGCs (Fig 3d). Consensus structural motifs in lanthiphage core peptide variants are described in Supplementary Fig 5c, with the motif TXDXCXXXCXXXC conserved in 57% of predicted core peptides. Since threonine, serine and cysteine residues that participate in lanthionine formation are conserved, such structural motifs might be key to their biological function.

### Diverse lanthiphages confer anti-phage defense

To further investigate and validate the anti-phage function of diverse lanthiphage types with different core peptide variants, we sought to express them in a heterologous host, which is devoid of any lanthiphage. We constructed *Streptomyces albus* J1074 strains for the constitutive heterologous expression of four different lanthiphages detected in *Streptomyces* genomes, each encoding distinct predicted core peptides (red stars in Fig 4a, Supplementary Fig 6). For each tested lanthiphage, we introduced all lanthiphage genes in synthetic operons to construct lanthiphage-expressing strains, and built corresponding control strains expressing incomplete BGCs (all enzymes except precursor peptide gene and LphE when applicable, Fig 4a, Supplementary Fig 6, Supplementary Table 3). The resulting lanthiphage-expressing strains were named lanthiphage_Sma, lanthiphage_Sc3, lanthiphage_Scl, lanthiphage_Sr2, according to their species of origin.

**Figure 4.**
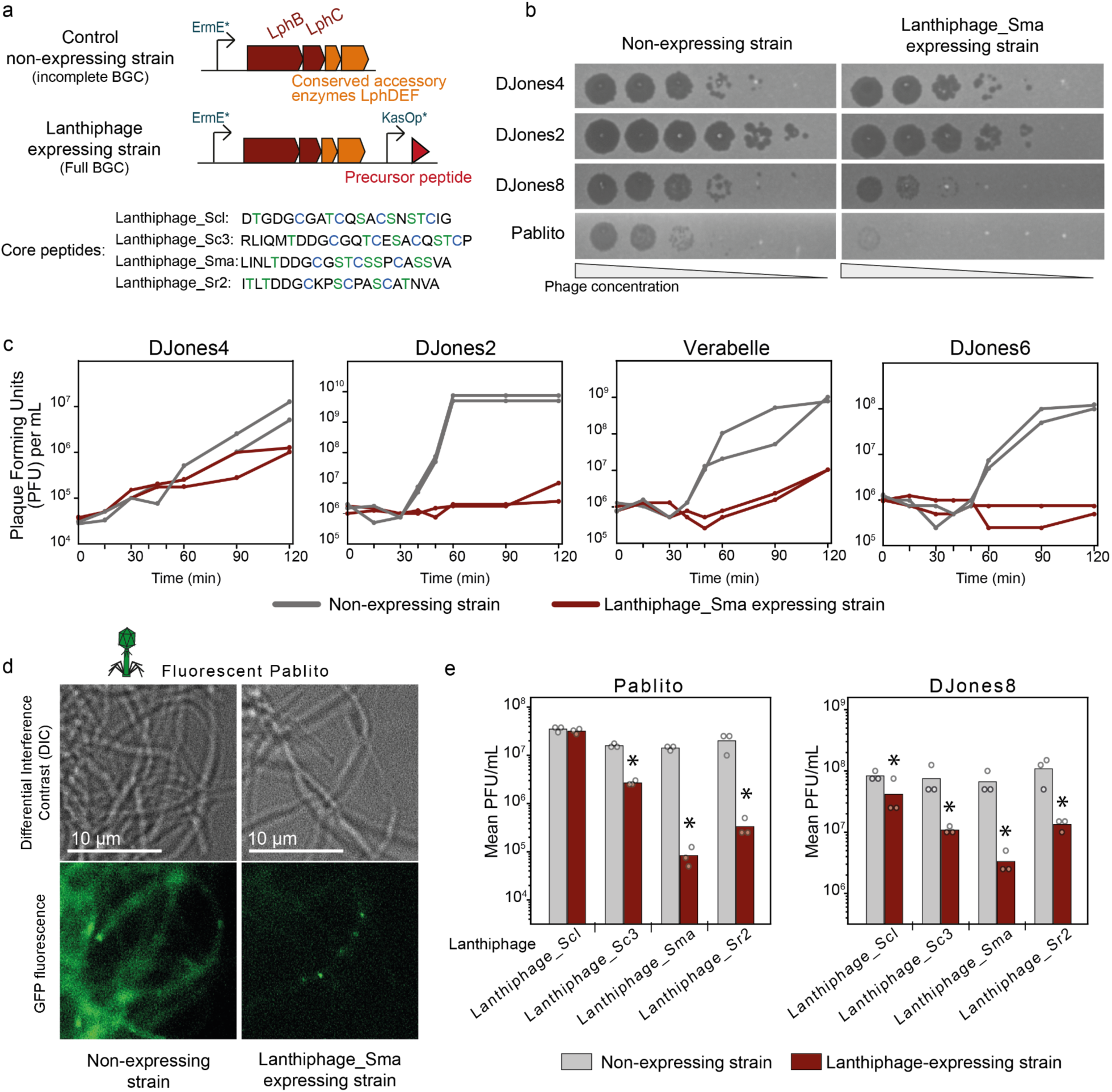
Heterologous expression of lanthiphages confers anti-phage defense. **a.** Schematic representation of synthetic operons designed to construct strains for assessing the anti-phage activity of lanthiphages. Lanthiphages cloned are Lanthiphage_Scl (clade 2, Type 2, core peptide variant 2), Lanthiphage_Sc3 (clade 5, Type 1, core peptide variant 2), Lanthiphage_Sma and Lanthiphage_Sr2 (both clade 9, Type 5, core peptides variant 6). A control strain expresses an operon encoding an incomplete lanthiphage under the control of the constitutive promoter ErmE*. The lanthiphage-expressing strain, includes an additional operon controlled by the strong constitutive promoter KasOp* expressing the remaining genes for full BGC expression. Detailed BGC refactoring is described in Supplementary Fig 6. **b.** Plaque assay results shown for four phages on strain expressing either the incomplete (control strain) or the full (lanthiphage-expressing strain) lanthiphage_Sma. Images are representative of three biological replicates. **c.** Liquid infection assays of control and lanthiphage-expressing strains infected at a multiplicity of infection (MOI) of 0.1. Phage replication was tracked by measuring plaque forming units (PFU) per mL of culture over time after infection. Two biological replicates are presented as individual curves. Additional curves for other lanthiphages and phages are in Supplementary Fig 8. **d.** Fluorescence microscopy pictures of cultures infected using a green fluorescent version of the phage Pablito at MOI of 0.1. Cultures were imaged 24 h after infection. Fluorescent dots in the lower right picture, correspond to auto-fluorescence of the septa in the hyphae observed in *Streptomyces* colonies. **e.** Efficiency of plating of phages Pablito and DJones8 on the four tested lanthiphage-expressing strains and their respective non-expressing control strain. Bars represent an average of three biological replicates, with individual data points overlaid. Stars indicate statistically significant reduction of PFU/mL compared to control strain (t-test, p-value<0.05). Results for all phages of the DJones collection are found in Supplementary Fig 7.

Next, we challenged lanthiphage-expressing strains with phages from the DJones collection. The lanthiphage_Sma expressing strain shows increased resistance against 9 of 11 phages, compared to the control non-expressing strain. For example, we observed that lanthiphage_Sma expression results in a 170-fold increase in resistance against phage Pablito^34^ in plaque assays, but has no effect on phage DJones4 (Fig 4b, Supplementary Fig 7). This trend was confirmed by liquid infection assays (Fig 4c, Supplementary Fig 8), where early inhibition of phages Verabelle, DJones2 and DJones 6 is observed in lanthiphage_Sma expressing cultures, but no inhibition of DJones4. Using fluorescence microscopy to track the replication of a fluorescent variant of phage Pablito (GFP-labeled major capsid protein), we confirm the inhibition of phage particles produced in the mycelium of lanthiphage_Sma expressing strains (Fig 4d). Finally, we demonstrate that all four tested lanthiphages provide clear anti-phage activity against at least one phage in plaque assays and liquid infections compared to control strains (Fig 4e, Supplementary Fig 7, Supplementary Fig 8). All phages from the DJones collection were inhibited by the expression of at least one tested lanthiphage, except DJones4 for which no effect was observed in all tested conditions. This indicates that lanthiphages provide protection against a subset of phages, implying a certain range of specificity in their anti-phage activity.

### Lanthiphage_Sco produces a novel lanthipeptide

To explore the mechanism of action of lanthiphages, we assessed the production of the lanthipeptide synthesized by lanthiphage_Sco under different growth conditions using untargeted metabolomics analyses of cell extracts by liquid chromatography-mass spectrometry (LC-MS). We identified a novel [M+2H]^2+^ ion m/z 1114.09 corresponding to a peptide mass of 2226.17 Da in extracts of lanthiphage_Sco expressing *S. coelicolor* as compared to non-expressing strains (Fig 5a, Supplementary Fig 9a,b). Knocked-down expression of lanthiphage_Sco genes (precursor peptide genes or LphB), resulted in a substantial reduction of this peptide’s signal in cell extracts (Supplementary Fig 9c) further confirming that lanthiphage_Sco is involved in its production. The observed molecular mass does not match any known lanthipeptide, and that of potential products predicted from the core peptide sequences of lanthiphage_Sco, suggesting additional unknown modifications, potentially catalyzed by LphDE. Tandem MS analysis revealed fragments from the C-terminal part of the core peptide and some related immonium ions (Supplementary Fig 9d), supporting that this peptide species is derived from lanthiphage_Sco. Complex MS2 patterns are consistent with the macrocyclized nature of the lanthipeptide (Supplementary Fig 9e). Overall, these results confirm the production of a novel lanthipeptide compound by lanthiphage_Sco of *S. coelicolor*.

**Figure 5.**
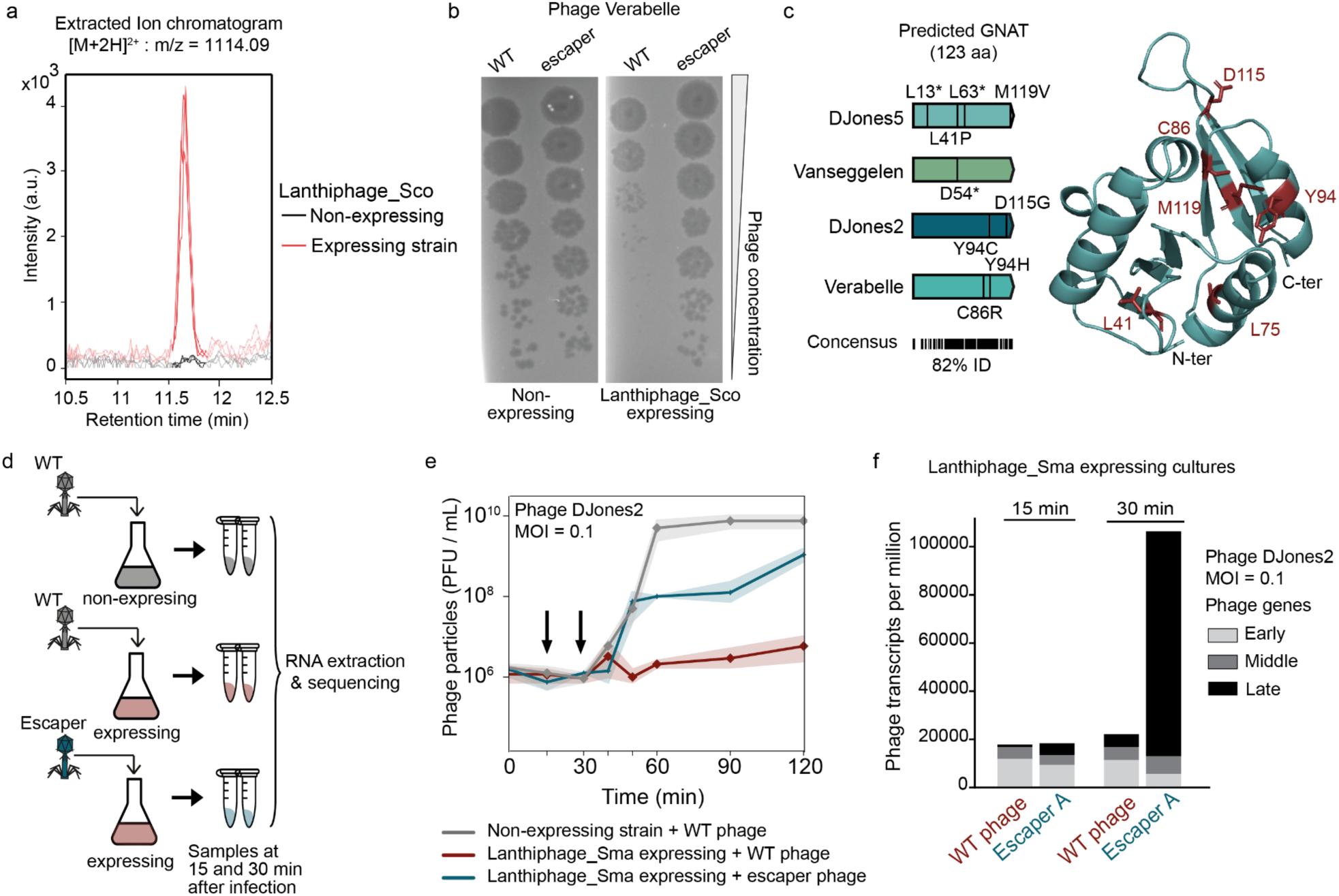
Lanthiphage_Sma alters phage transcription. **a.** Representative extracted ion chromatograms of a [M+2H]2+ ion corresponding to a peptide mass of 2227.2 Da present in extracts from the lanthiphage_Sco expressing strain, and absent in extracts from the non-expressing control strain, grown in solid SFM medium. Signals from three biological replicates are presented as individual curves. **b.** Plaque assay results shown for wild-type phage Verabelle and escaper A on lanthiphage_Sco expressing compared to the non-expressing *S. coelicolor* strain. Images are representative of three biological replicates. **c.** Location of representative mutations found in phages that escape lanthiphage defense. All isolated escaper phages harbor a mutation in a gene encoding a predicted GNAT, with a majority of single-point mutations that result in a single amino acid mutation or a premature stop codon. Single amino acid mutations are highlighted in red on the AlphaFold model of the GNAT from phage DJones2. All mutations and escaper variants are described in Supplementary Table 4. **d.** Schematic representation of liquid infection assays performed to assess effect of lanthiphages on phage transcription. Lanthiphage-expressing and non-expressing strains were infected with the WT or escaper A variant of DJones2 at a MOI of 0.1. **e.** Phage plaque forming units (PFU) per mL of cultures was measured over time after infection. Curves represent the mean of three biological replicates, and the shaded area corresponds to the 95% confidence interval. Black arrows indicate time point for sample collection. Additional curves for other lanthiphages and escaper phages are in Supplementary Fig 8. **f.** Level of transcription of phage genes at different time points after infection of lanthiphage-expressing strains. Bars represent the mean transcripts per million of two biological replicates, and gray stacks represent the proportion of early, middle and late phage genes transcribed (see Methods).

### Lanthiphage_Sma alters phage transcription

To determine whether the lanthipeptide compounds produced by lanthiphage-expressing strains are secreted and can protect susceptible non-expressing strains, we analyzed the anti-phage activity of cell extracts and spent media of recombinant lanthiphage-expressing *S. albus* cultures and their controls. No significant effect on phage replication or cell viability was observed from cell extracts or spent media from lanthiphage-expressing cultures compared those from control strains (Supplementary Fig 10a). While these results cannot fully rule out lanthipeptide secretion, they would be consistent with an intracellular action of the compounds produced by lanthiphages. This hypothesis is further supported by the absence of ABC transporter genes usually involved in lanthipeptide secretion^28^ near the 4 lanthiphages tested, in line with observation that only a small fraction of all lanthiphages are located next to genes for ABC transporters (18.3% versus 63,4% in other class I lanthipeptide BGCs (Supplementary Fig 1c,e).

As many characterized lanthipeptides have antibacterial activity^35^, we tested whether lanthiphage expression affects the cell viability. We compared the growth, fitness and development of lanthiphage expressing and non-expressing strains, and observed no significant differences (Supplementary Fig 10b,c,d). Since previously characterized lanthipeptides interact with cell wall and membrane components^35^, we hypothesized they might block phage adsorption. However, phage adsorption assays, which measure the ability of phages to attach to bacterial cells, show that lanthiphage expression does not affect phage adsorption compared to control strains (Supplementary Fig 10e). Overall, these results further support an intracellular mechanism of action following phage adsorption, directly targeting the phage cycle without being toxic to the host.

To further shed light on the effect of lanthiphage-produced compounds on phage replication, we isolated multiple escaper mutants^36^ of four different susceptible phages from the DJones collection (DJones2, DJones5, Vanseggelen, Verabelle) on lanthiphage-expressing *S. coelicolor* strain, for which we observed the highest anti-phage activity (Fig. 5b, Supplementary Fig 11a). These phage mutants also escape the effect of heterologous lanthiphage expression in recombinant *S. albus* strains (Supplementary Fig 11b), suggesting a common anti-phage mechanism. Genome sequencing of 15 isolated escaper phages (from four different phages) shows that all have mutations in the homolog of the same gene which codes for an uncharacterized Gcn5-related N-acetyltransferase (GNAT) (four homologs that share a minimum of 82% protein sequence identity between each other) (Fig. 5c). Most mutations (12/15) consist of single-point mutations resulting in single amino acid changes or a premature stop codon (Fig. 5c, Supplementary Table 4). Recent studies have shown that phage-encoded GNATs can regulate phage gene transcription, by controlling the shift between early and late gene expression^37^. While the predicted GNAT does not resemble previously described enzymes, we hypothesized it could impact transcriptional regulation. To test this hypothesis, we analyzed the host and phage transcriptional levels during early infection of lanthiphage_Sma expressing strains by phage DJones2 WT or escaper A (Y94C mutant) (Fig. 5d,e). While the host transcriptional program is similar during infection by WT and escaper phage (Supplementary Fig 12a), the transcriptional program of the WT phage is altered by lanthiphage expression (Fig. 5f, Supplementary Fig 12b,c). At 15 minutes post-infection, the number of transcripts is comparable between both WT and escaper phages (15k per millions). However, at 30 min post-infection, WT phage transcripts were five times less abundant compared to escaper phage (Fig. 5f). When analyzing the differentially expressed genes, we observed 15 times fewer transcripts of late genes in WT phages compared to escaper phages (Fig. 5f, Supplementary Fig 12b). These results indicate that lanthiphage expression inhibits global phage transcription and specifically alters the expression of late genes of the WT phage compared to the escaper phage. Quantitative PCR (qPCR) analysis of injected phage DNA shows that these differences in transcription levels occur with similar amounts of injected phage DNA in the host (Supplementary Fig 12d). Our results suggest that the compounds produced by lanthiphages affect phage transcription through a mechanism dependent on a phage-encoded GNAT.

## Discussion

Our work describes the discovery of a diverse family of >2000 lanthipeptide BGCs with anti-phage function, called lanthiphages, revealing that RiPPs can participate in anti-phage defense. We demonstrate the anti-phage function of five different lanthiphages in native or heterologous hosts. Lanthiphages are widespread in Actinobacteria but so far restricted to this phylum, which suggests that they may be adapted to the specific metabolism, multicellular lifestyle and ecological niches of this lineage. Our genomics analyses reveal a single evolutionary origin of lanthiphages from a non defensive clade of class I lanthipeptides. We uncovered unique features of lanthiphages such as the presence of specialized genes *lphDEF* and a shared motif in the core peptides while demonstrating the diversity of the family which we classified in seven major types. Finally, we show that lanthiphages act intracellularly by targeting phage transcription.

Beyond our initial findings, many aspects of lanthiphage biology remain unexplored. Our study only examined a limited subset of lanthiphages, and other types require experimental characterization. Key questions persist regarding the chemical structure of lanthipeptides produced by lanthiphages, their diversity, and structure-to-function relationships. For example, using similar LC-MS methods on lanthiphage-expressing *S. albus* strains, we could not detect the presence of a specific peptide compared to controls, hence requiring more research to determine the mass of the lanthipeptides produced. The specific activity of each enzyme within these BGCs also needs further investigation. Additionally, the exact mechanisms through which lanthiphages exert their anti-phage effects are still unclear. For example, it is possible that lanthiphage-produced compounds interact with the identified phage GNAT, preventing the transcriptional shift between early and late phage genes, but this hypothesis requires further study.

While much remains to be understood about lanthiphages, the discovery of this novel family marks a significant expansion in the number of BGCs involved in chemical defense. Our study highlights that the emergence of BGCs linked to anti-phage defense can be identified genomically through associations with defense islands. Specifically, we demonstrate a previously unknown biological role of lanthipeptides in protecting bacteria from phage infection. This discovery raises the possibility that other types of BGCs might also be involved in anti-phage defense. We hypothesize that additional anti-phage BGCs could be predicted using a similar genomic approach to the one we employed.

Unveiling the function of natural products has traditionally been challenging, often requiring compound isolation or random screening assays. Our study presents a novel method that employs genomic clues to predict the anti-phage activity of BGCs. Whether other genomic clues could be used to predict other functions remains to be determined. Furthermore, natural products have historically been a fruitful source of clinical drugs, such as antibiotics. Our research reveals new anti-phage RiPPs. Given the recently identified conservation of anti-phage defense mechanisms between bacteria and humans, this discovery hints that bacterial natural products may offer a reservoir of novel antiviral compounds potentially usable against human pathogens.

## Materials and methods

### Genomic detection of lanthipeptide BGCs near known defense systems and within mobile genetic elements

Genomic detection of lanthipeptide BGCs was performed using antiSMASH v6.1.1^25^ on the 29,414 genomes of Actinomycetes from the RefSeq database (all assembly qualities: complete, chromosome, scaffold and contigs). The core biosynthetic genes of each BGC were retrieved using the antiSMASH annotations (precursor lanthipeptide, dehydratase, and lanthionine cyclase genes). Anti-phage defense systems were detected using DefenseFinder^26^ v1.2.1 with models v1.2.3. To determine the proximity of lanthipeptide BGCs of all classes to known defense systems, we assessed the presence of a defense system within 23 genes upstream or downstream of the lanthionine cyclase of each BGC (20 genes + 3 genes for the rest of the lanthipeptide BGC). The random protein defensiveness statistic was computed by sampling 10,000 proteins using shuf. The defensiveness statistic was calculated with the same method as for the lanthionine synthetase. All lanthionine cyclase sequences were then clustered at 80% identity and 80% of coverage using MMseqs2^38^ v13.45111. For each cluster, we computed a defense score which corresponds to the proportion of cluster members that have at least one defense system in their vicinity. To compute a defense score at the lanthipeptide class level (lanthiphage, non-defensive class I lanthipeptide, class I-IV), we took the average score of all clusters belonging to that class where all the clusters have the same weight. Prophages and plasmids were detected using genomad^39^ v1.7.4. A lanthipeptide was considered to be inside a prophage if the cyclase was present inside the prophage boundary +-10kb. A lanthipeptide BGC was considered to be inside a plasmid if it was present inside a replicon detected as plasmid by genomad.

### Detection and filtering of lanthionine cyclases

A total of 14,437 synthetase sequences were initially retrieved using the AntiSMASH tool. These sequences were then filtered based on their size, resulting in 13,815 sequences (size thresholds for each class: I: 300-550 amino acids, II: 800-1200 amino acids, III: 800-950 amino acids, and IV: 800-1000 amino acids). Lanthionine cyclases were clustered using MMseqs2 with sequences identity set at 80% and coverage set at 80%. This clustering process yielded 3,220 representative sequences, distributed as follows: Class I: 1,180 sequences, Class II: 703 sequences, Class III: 903 sequences, and Class IV: 434 sequences.

### Phylogenetic tree construction

For all the different phylogenetic trees, sequences were first aligned with muscle super5 v5.1^40^ and trimmed using clipkit v1.3.0. Phylogenetic tree construction was performed using IQ-TREE v 2.2.3 with models finder and 2000 ultrafast bootstrap. All multiple sequence alignments and phylogenetic trees are available on Github (See Data Availability).

### Phylogenetic tree sequences selection

The protein tree of the LANC_like domain of all lanthionine cyclase clades (Supplementary figure 1b), was built using all previously detected 3220 representative sequences (from all classes) as well as MiBIG BGC lanthionine synthetase. The MiBIG v3.1 was downloaded https://dl.secondarymetabolites.org/mibig/mibig_gbk_3.1.tar.gz on May 19 2024. Lanthionine synthetase from the MiBIG BGC were retrieved using AntiSMASH annotation. First, we extracted the LANC_like domain (PF05147) from the 3,284 proteins. To do so, an alignment was built for each class of lanthipeptides using MAFFT. This allowed us to extract the LANC_like_domain with confidence thanks to good quality alignments. The start and end positions of the LANC_like domain (PF05147) were identified on each of these alignments using HMMER’s hmmsearch^41^. Once the domains were retrieved, they were realigned all together following the procedure described above (see **Phylogenetic tree construction**). The class I lanthionine cyclase tree (Figure 1d, Supplementary Figure 1c) was computed using n = 1,180 representative sequences of the class I lanthionine cyclase. To root the tree a n = 30 outgroup was created by a random sampling of previously detected class II LANC_like domains (sequences available on Github). For the lanthiphage tree (Figure 3a, Supplementary Figure 5a), all homologs of LphC were used (n = 2063). To root the tree n = 20 non-lanthiphage class-I lanthionine cyclase were manually selected (sequences available on Github).

### Strains and growth medium

Bacterial strains and plasmids used in this work are listed in Supplementary Table 5. *Streptomyces* strains were routinely maintained on Soya-Flour-Mannitol (SFM) agar medium, supplemented with the antibiotics apramycin (50 µg/mL) and hygromycin (100 µg/mL) when appropriate. Experiments to study phage-bacteria interactions were done in Difco Nutrient Broth (DNB) medium supplemented with 0.5% glucose and 4 mM Ca(NO_3_), or Maltose Yeast Extract (MYM) medium (4 g/L maltose, 4 g/L yeast extract, 10 g/L malt extract).

### Bacteriophage isolation, propagation and sequencing

The isolation of bacteriophages from soil samples of different locations was carried out following the “enrichment” phage isolation protocol previously described^34^. Soil samples were collected in France (as part of the citizen science project ‘Science à la Pelle’^42^ and in the Netherlands (Leiden University). Briefly, soil samples of each location, 0.5 gram each, were added to separate 50 ml Falcon tubes containing 10 ml of DNB medium or MYM medium. To each tube, 5 µl of each spore stock of *S. coelicolor*, *S. griseus*, *S. venezuelae* or *S. lividans* were added and cultures were incubated overnight with shaking at 30°C. Cultures were then centrifuged at 4000 x g for 10 minutes, and the supernatants were filter sterilized and stored in the dark at 4°C until further use. Isolation of single phages was carried out by plating serial dilutions of filtered supernatants on DNB soft agar plates, and inoculated with approximately 10^8^ spores of the bacterial host that was used for propagation. Plates were incubated for 24-48 hours at 30°C until plaques appeared. High titer phage stocks from single plaques were obtained by picking each plaque and introducing them to 10 mL DNB broth inoculated with 10 µl of *S. albus* spores, and incubated overnight at 30°C with shaking. Lysates were then filtered through a 0.45 µm Millipore filter into sterile 15 ml tubes and individual viruses were stored in the dark at 4°C until further use. Consequently for each isolated phage, lysate stocks were routinely maintained by amplification using *S. albus* in DNB medium. DNA isolation of phages from cell lysates was performed using the Phage DNA Isolation Kit (Norgen Biotek Corp.) and quantified with a Qubit dsDNA Assay kit. Whole genome sequencing was performed by the Institut Pasteur Biomics Platform and a *de novo assembly of the* phage genomes was performed using SPAdes v3.15.5. The genomes of these phages were characterized (Supplementary Fig 2), and belong to the family of Arquatrovirinae. We named the collection after Doris Jones Ralston (1921-2011), a microbiologist who discovered anti-phage natural products from Actinobacteria^9,10^. This collection is composed of the previously described phages Pablito^34^, Verabelle and Vanseggelen^43^ and novel phages listed in Supplementary Table 5.

### Plaque assays

Square plates containing 35 mL of either DNB soft agar (0.5% agarose (w/v)) or MYM soft agar (0.5% agarose (w/v)) were inoculated with approximately 10^8^ spores of the strain of interest. Plates were dried and 4 µL droplets of serial dilutions of each phage were placed on the agar surface, with dilutions ranging from 10^-1^ to 10^-7^. Plates were incubated at 30°C overnight and placed at room temperature for another 24h, and plaque forming units were counted to evaluate phage replication.

### Knocking-down lanthipeptide producing genes

For knocking down genes involved in lanthiphage production, we used the engineered Cumate-Based Inducible CRISPR interference (CUBIC) system^32^ in *S. coelicolor*. Protospacers sequences targeting the precursor peptide genes and serine/threonine dehydratase gene of the lanthiphage BGC from *S. coelicolor* M145 (NZ_CP042324.1) were designed using the ‘Find CRISPR’ tool of the Geneious Prime software (version 2023.2.1). The protospacer sequences were synthesized as oligos (found in Supplementary Table 6) and integrated in the vector pCB-1 by Golden Gate Assembly. Plasmids were delivered in *S. coelicolor* by intergeneric conjugation according to standard protocols^44^ in SFM medium supplemented with 10 mM MgCl_2_ and hygromycin. Three resulting conjugants were picked and spore stocks were prepared on SFM plates supplemented with cumate (20 µM). Resulting vectors and strains are listed in Supplementary Table 5. To test the anti-phage resistance of knock-down strains, plaque assays were performed as previously described in DNB soft agar with 20 µM cumate.

### Detection of conservon systems near lanthipeptide BGCs

To detect conservon systems we used MacSyFinder^45^ v2.1.1 with a coverage threshold of 0.4 and custom MacSymodels of the conservon system (Available on Git link). The HMM protein models of the conservon proteins were built using BLASTp^46^ results (50% identity 80% coverage) of *Streptomyces coelicolor* M145 (NZ_CP042324.1) Cnv8 proteins (Cnv8A, Cnv8B, Cnv8C, Cnv8D, CnvF8) on the RefSeq nr database. Proteins were aligned using MAFFT^47^ v7.505 (auto mode) and HMM profiles were created using HMMbuild^48^ (HMMER v3.3.2) and a GA threshold of 20 was applied to all profiles. The detection definition was to detect at least 2 genes to be detected as a conservation system.

### Detection of Lanthiphage on the Prokaryotic RefSeq database

Genomes from the RefSeq complete genomes (n = 22,803) downloaded in July 2022 were used to determine if the lanthipeptide was present in other phyla beyond Actinomycetota. Lanthiphage were detected using custom MacsyFinder models of the lanthiphage defense systems (available on Github). This model is composed of 19 HMM profiles that were built using the results of the antiSMASH detection.

### Detection of domains present in Lanthiphage BGCs

The different domains present in lanthipeptide class I were detected using antiSMASH^25^ results. The domains detected by antiSMASH were LANC_like (PF05147), Lant_dehydr_C (PF14028), Lant_dehydr_N (PF04738), PCMT (PF01135). Transporter proteins were detected using the antismash annotation where the “gene_function” is a transporter. The hypothetical protein present in several clades of lanthiphage was detected using custom HMM profiles available on the GitHub repository (https://github.com/mdmparis/lanthiphage_2024).

### Core Peptide sequence classification

The tool antiSMASH v6.1.1 was used to define the sequences of the predicted core and leader peptides from the sequence of the precursor lanthipeptide of a given BGC. Precursors where antiSMASH did not split between leader and core peptide were compared using BLASTp against core peptide described by Walter et al.^30^. For each of those precursors, the best hit against the database was used to delimit the potential boundary between the leader and the core peptide. All the core peptides (antiSMASH annotated and manually delimited) were then compared using BLASTp against each other. A proximity score was calculated (the % identity divided by the % coverage). We used this proximity score to compute a distance matrix that was clustered with Louvains methods using python-louvain^49^ v0.16. In total, we defined 9 variants, with more than 10 representatives. The sequence logo for each variant, and for all core peptides from the defensive and non-defensive clades were generated using WebLogo^50^ 3.7.12.

### Construction of *Streptomyces* strains for expression of defensive lanthipeptide BGCs

The sequences of the lanthiphage BGCs identified in the genomes NZ_CP015726.1 (*Streptomyces sp. RTd22*), NZ_JAIY01000006.1 (*Streptomyces sclerotialus*), NZ_LJIW01000001.1 (*Streptomyces malaysiensis*), NZ_FOLM01000022.1 (*Streptomyces sp. CNT318*) were retrieved. The original coding sequences for each BGC were refactored into 2 different synthetic operons, referred to as operon A and operon B (Supplementary Fig. 6). For operon A: all core modifying enzymes genes (except for the predicted FxlM methyltransferase) predicted by antiSMASH were kept in their natural chromosomal organization, a synthetic RBS sequence was placed before the START codon of the first enzyme, and the operon was put under the control of the constitutive ErmE* promoter. When applicable, a conserved hypothetical protein of unknown function was included when present in the native operon. For operon B: the precursor lanthipeptide gene was placed under the control of the constitutive KasOp* promoter and a synthetic RBS before its START codon; when applicable the predicted FxlM methyltransferase present in the BGC was placed after the lanthipeptide precursor gene on this same operon, with a synthetic RBS sequence. For each BGC to test, 2 vectors were cloned: one vector expressing the full lanthipeptide BGC (operons A and B), one negative control vector expressing an incomplete BGC (operon A only). The sequences of the designed operons were synthesized by Genscript and cloned into the pOJ436 cosmid vector^51^. Operons A were cloned into the vector at the PvuII restriction site of pOJ436, and operins B into the BglII site. These constructs were then introduced in the chromosome of the strain *S. albus J1074* by conjugation from *Escherichia coli* S17 following standard protocols^44^. Three positive *S. albus* conjugants were screened on a growth medium supplemented with apramycin and the chromosomal integration of the BGCs was confirmed by colony PCR using the primers listed in Supplementary Table 6.

### Liquid infection assays

Liquid cultures were done in Erlenmeyer flasks with DNB media inoculated with spores of interest at OD_450_ = 0.15. Cultures were incubated for 5h with shaking at 30°C, then phages were added at a multiplicity of infection (MOI) of 0.1. After infection, 1 mL of culture was sampled every 15 minutes (unless otherwise noted), immediately filter-sterilized, and stored at 4°C until use. We then assessed the number of phage particles in each sample by titrating them on DNB soft agar medium inoculated with WT *S. albus J1074* spores. Plates were incubated at 30°C overnight and placed at room temperature for another 24h. Plaque forming units were counted to evaluate phage replication.

### Construction of fluorescent phage Pablito

To construct a fluorescent version of phage Pablito, we constructed the integrative vector pGWS1803 (derived from pSET152^51^ (Bierman, Logan et al. 1992)), which carries the gene encoding the Pablito capsid protein, fused to *egfp* and controlled by the *gap* promoter. The *gap* promoter was amplified from *S. coelicolor* M145 genomic DNA using primers Pgap_F and Pgap_R. The coding sequence for phage Pablito capsid protein was amplified from genomic DNA of MBT86 Pablito revertant using primers Pablito_CP_F and Pablito_CP_R. The *egfp* gene was amplified from pGWS785^52^ with primers 1572_P3 and 1572_P4. The sequence of primers used are in Supplementary Table 6. These fragments were then cloned into EcoRI and XbaI digested pSET152 via Gibson assembly to generate pGWS1803. The resulting construct, pGWS1803, was subsequently conjugated into MBT86 Pablito revertant as described previously^44^. The spores of the generated exconjugants were inoculated in 10 mL of Tryptic Soy Broth with Sucrose (TSBS) and incubated overnight at 30°C with shaking. A 100 µL aliquot of this overnight culture was then transferred to 10 mL DNB containing 0.5 µg/ mL of mitomycin C and incubated again overnight under the same conditions. After incubation, the culture was filtered through a 0.22 µm filter. The filtered culture was then used for a lytic plaque assay on MBT86 strain. The plaque assay plate was incubated at 30°C for one day until plaques appeared. Subsequently, 20 mL of DNB was added to the plaque assay plate, and the plate was gently shaken for 2 hours to recover the phages. The DNB broth containing the fluorescent phage Pablito was then filtered again through a 0.22 µm filter to obtain a single high-titer phage stock.

### Microscopy

To image the infection of fluorescent phage Pablito (gift from Leiden University) in the strains LP_Malay_full and LP_Malay_neg, 10^6^ spores were inoculated in 10 mL DNB medium, infected with fluorescent Pablito at MOI=0.1 and incubated overnight at 30°C with shaking. After 24 hours, confocal images were made using a Zeiss LSM900 Airyscan 2 microscope. The fluorescent phage was excited at 488 nm and monitored at 509 nm. To assess the morphology and development of *Streptomyces* strains, our engineered strains were grown in DNB for 24 hours at 30°C with shaking, and brightfield images were made with a Zeiss Axio Lab A1 upright microscope, equipped with an Axiocam 105 color camera. Subsequently, hyphal thickness (n=6) was measured with ImageJ. All microscopy images were processed using OMERO.

### Biomass assay

To measure the biomass of engineered strains after phage infection, 10^6^ spores of the strain overexpressing the lanthiphage BGC from *Streptomyces sp.* RTd22, or its corresponding negative control strains, were inoculated in 10 mL DNB medium at 30°C with shaking. After 5 hours of germination, phage Pablito or phage DJones3 was added at MOI=0.3. After overnight infection, the remaining biomass was recovered using a 5 µm pluriSelect cell strainer, dried and weighed.

### Lanthipeptide extraction and LC-MS/MS analysis

Spores of the recombinant *Streptomyces* strains were spread on solid media (SFM or MYM) and left to grow at 30°C for 7 days. A portion of the mycelium and the media was taken using the wide extremity of a 1 mL tip and extracted with 1 mL methanol at room temperature for 4 hours. The extracts were dried by speedvac and re-suspended in 200 µL 80% acetonitrile (ACN). Ultra-high-performance liquid chromatography-MS analysis was performed on an Ultimate 3000-RSLC system (Thermo Scientific) connected to an electrospray ionization-quadrupole–time of flight instrument (Maxis II ETD, Bruker Daltonics). One µL of the extract was analyzed on a RSLC Polar Advantage II Acclaim column (2.2 μm, 120 Å, 2.1 × 100 mm, Thermo Scientific) with a linear gradient (2-80 % in 15 min) of mobile phases formic acid (FA) 0.1% and LC-MS grade ACN + FA 0.08%, at 300 μL/min. The LC-MS data were collected in positive mode in the range m/z 300 - 1600, using collision-induced dissociation in data dependent auto-MS/MS mode. Targeted MS/MS were performed on the ions of interest using the MRM mode. The MS instrument was calibrated at each run start using a sodium formate solution consisting of 10 mM sodium hydroxide in isopropanol / 0.2% formic acid (1:1 v/v). The LC-MS and LC-MS/MS data were analyzed and converted to NetCDF and mgf format using DataAnalysis software (version 4.4, Bruker Daltonics)

### Colony morphology

To determine if there are any differences in colony morphology between the negative control and lanthipeptide strains, 10^5^ spores were spotted on MYM plates. After 7 days of incubation at 30°C, images of the edge of the colonies were made with a Zeiss SteREO Discovery.V8 stereomicroscope equipped with a Bresser MikroCam SP 5.0.

### Assessing anti-phage activity of spent media and cell extracts

To assess the effect of spent media from lanthiphage expressing cultures, 5mL of DNB media and supplemented apramycin (50µg/mL) were inoculated with spores from lanthiphage expressing and non-expressing strains, and incubated 72h at 30°C with shaking. 1 mL of these cultures was used to inoculate 25 mL of DNB media in Erlenmeyer flasks. After two days of incubation at 30°C with shaking, the cultures were centrifuged, and the supernatant was filtered. New cultures were inoculated in 4.5 mL of filtered supernatant and 500 µL of 10X DNB with 10^6^ WT *S.albus* spores, and with or without 10^5^ phages (MOI of 0.1). Cultures were incubated overnight at 30°C with shaking. CFUs of non-infected cultures were calculated by spotting serial dilutions of cultures onto solid MYM, and PFU/ mL of infected cultures was calculated as previously described.

To assess the anti-phage effect of cell extracts from expressing and non-expressing strains (prepared as previously described), 2 μL of cell extracts were added to 250 µL DNB inoculated with 10^6^ WT *S.albus* spores and 10^5^ phages (MOI of 0.1). Cultures were incubated at 30°C with shaking, and samples were taken 1 h, 2 h and 24 h after infection to quantify PFU/mL.

### Absorption assays

Liquid cultures of 20 mL DNB medium were inoculated with 10^7^ spores of strain of interest in an Erlenmeyer flask. Cultures were incubated at 30°C for 5h with shaking. Phages were then added at an MOI of 0.1. After infection, 100 µL samples of cultures were taken every 10 min until 1h of incubation and transferred into an Eppendorf tube containing 900 µL DNB medium. Samples were vortexed and centrifuged for 7 min at maximum speed. Supernatants were filtered, sterilized and stored at 4°C. Phage titers were determined as previously described.

### Isolation of phage escapees

Each phage lysate was diluted and spread on the surface of soft DNB agar plates inoculated with spores of the strains *S. coelicolor ΔcvnF8*, to get single lysis plaques. Each lysis plaque was picked and amplified in DNB media inoculated with WT *S. albus* spores, by incubation at 30°C overnight with shaking. Filter sterilized lysates were used to compare the sensitivity of the amplified phages to the WT phage, by performing plaque assays on plates of WT *S. coelicolor* and *S. coelicolor ΔcvnF8*. Phages were considered as escapees when their infection was not inhibited by the lanthipeptide, as compared to the WT phage. The collected escapees were then amplified and sequenced as previously described.

### Transcriptomics via RNA sequencing

To compare transcription of the host strains and phage DNA during infection with phage DJones2 and the escaper variant A, we performed liquid infection assays in Erlenmeyer flasks with DNB media inoculated with spores of interest at OD450 of 0.15. Cultures were incubated for 5h with shaking at 30°C, then the phage of interest was added at a multiplicity of infection (MOI) of 0.1. At 15 min and 30 min after infection a total of 3 OD units of cells were harvested by centrifugation at 5,000 × g and 4°C for 10 min. Cell pellets were washed twice with phosphate-buffered saline (PBS) and stored at −20°C. RNA purification, depletion of rRNA, library preparation, and sequencing were conducted by Genewiz (Leipzig, Germany).

Transcriptomics data was analyzed as follows. Reads from the RNA sequencing were trimmed using trim galore (https://github.com/FelixKrueger/TrimGalore) v0.6.10 with a quality threshold of 25 and minimal length of 60 bp. RNA reads quantification was performed using salmon53 v1.10.2 resulting in both raw read counts per genes and transcript per million. Read count fold changes depending on condition were calculated using PyDESeq254 v0.4.7. Early, middle and late phage genes were characterized using the ratio of the normalized transcript per million between 15 and 30 minutes in the negative control strain. For the differential expression representation (Supplementary Figure 12a), the maximum LogFold change for visualization was set to six and the minimal adjusted p value was set to 10^-30^.

### Quantitative PCR (qPCR)

Culture conditions and cell pellets were prepared as described above in “Transcriptomics via RNA sequencing”. At 15 min and 30 min after infection a total of 3 OD units of cells were harvested by centrifugation at 5,000 × g and 4°C for 10 min. Cell pellets were resuspended in 500 µL PBS, mixed with Lysing matrix B (MP Biomedicals) and disrupted mechanically using a FastPrep-24 bead-beater device (MP Biomedicals) (2 cycles of 40 s at 4 °C). Cell lysates were then centrifuged at 12,000g for 10 min at 4 °C. Resulting supernatants were placed in clean tubes and DNA concentrations adjusted to 10 ng/μL using Nanodrop measurements. Diluted supernatants were used as the template DNA (5 μL) for qPCR using a 2×Luna universal qPCR master mix (New England BioLabs), with 0.5 μM of primers. Primers were designed following New England BioLabs guidelines to target the amplification of a phage gene coding for a hypothetical protein of DJones2 (ID: Djones2_00013), and a housekeeping gene of *S. albus* coding a DNA gyrase subunit B (ID: AGI89357.1). Measurements were performed using a Bio-Rad CFX96 C1000.

## Acknowledgements

We are grateful to Matthew F. Traxler (UC Berkeley) for the gift of conservon knock-out strains, and Vincent Libis (INSERM U1284) for the gift of the strain *S. albus J1074* and the pOJ436 vector. We thank Amel Abdennour, Uarda Kullolli, Katell Menard and all the participants of the project ‘Science à la Pelle’ for their contribution in the construction of the phage collection. We thank Marc Monot and Laurence Ma, from Biomics Platform (C2RT, Institut Pasteur, Paris) supported by France Génomique (ANR-10-INBS-09) and IBISA, which performed phage DNA sequencing. Several bioinformatic analyses were performed on the Core Cluster of the Institut Français de Bioinformatique (IFB) (ANR-11-INBS-0013). The LC-MS data were acquired at the MNHN Bioorganic Mass Spectrometry platform. We would like to thank Mehdi Beniddir, Pedro Leao, Raquel Castelo Branco, Somayah Elsayed, Hien Le for fruitful discussions and feedback, as well as Erick Denamur and the members of INSERM U1137 for their support in the early stages of this work. We are grateful to members of the MDM lab, as well as Beatriz Beamud, Enzo Poirier, Vincent Libis, Ariel Lindner and Flora Vincent for their useful comments on earlier versions of the manuscript.

## Authors contributions

H.S and A.B conceptualized the project. H.S. and F.T. carried out bioinformatic and phylogenetic analyses of lanthiphages, with early assistance from E.M.; H.S. designed and constructed strains for lanthiphage expression; H.S., M.G. and V.O. performed experiments to assess anti-phage activity and mechanism of action of lanthiphages; M.G. constructed CUBIC strains and performed related experiments; H.S., Y.L. and S.Z. performed LC-MS experiments and analyzed related data; H.S., V.O., M.L.B., H.G., and D.R. built phage collection; L.Z. constructed the fluorescent phage; H.S. and M.G. isolated and characterized phage escapers; F.T. analyzed DNAseq and RNAseq data; A.B. supervised the project. H.S., F.T., M.G. and A.B. wrote the initial version of the manuscript. All authors contributed to the design of the experiments, the discussion of the results and the final version of the manuscript.

## Data availability

All the data, including phylogenetic trees, HMM profiles and protein structure predictions generated with AlphaFold2, are available at https://github.com/mdmparis/lanthiphage_2024.

