## Supplementary Figures for "A family of lanthipeptides with anti-phage function"

### List of Supplementary Figures and Tables

**Supplementary Figures**

Supplementary Figure 1 – Genomic analysis of defensive clade
Supplementary Figure 2 – Doris Jones phage collection
Supplementary Figure 3 – Effect of lanthiphage_Sco expression on anti-phage resistance
Supplementary Figure 4 – Distribution of lanthiphages across Actinobacteria
Supplementary Figure 5 – Lanthiphages tree, genetic organization and core peptide
Supplementary Figure 6 – Design of synthetic operons for lanthiphage expression
Supplementary Figure 7 – Plaque assays of heterologous lanthiphage-expressing strains
Supplementary Figure 8 – Liquid assays of l heterologous lanthiphage-expressing strains
Supplementary Figure 9 – LC-MS characterization of compound produced by lanthiphage_Sco
Supplementary Figure 10 – Effects of lanthiphage expression
Supplementary Figure 11 – Characterization of lanthiphage phage escapers
Supplementary Figure 12 – Lanthiphage effect on host and phage transcription

**Supplementary Tables**

Supplementary Table 1 – List of analyzed class I lanthipeptide BGCs
Supplementary Table 2 – List of lanthiphage precursor peptides
Supplementary Table 3 – Sequence of synthetized operons
Supplementary Table 4 – Escaper phages
Supplementary Table 5 – Strains and phages used in this study
Supplementary Table 6 – Oligonucleotides used in this study

### Supplementary Text 1

#### Doris Jones Ralston

We named the collection of phages after Dr. Doris Jones (1921- 2011), a microbiologist from the United States of America. Her first works during her master's degree in the Waksman lab at Rutgers University focused on chicken microbiota to investigate the role of microorganisms and antibiotics in viral infection[1] contributing to the discovery of one of the first strains of *Streptomyces griseus* producing streptomycin[2]. She went on to work on bacteriophages and described the production of anti-phage substances, notably from actinomycetes[3]-[5] She earned her PhD from Berkeley University where she further worked as an immunologist.
To learn more: https://asm.org/articles/2024/march/women-in-the-history-of-antimicrobial-development

1. Jones, Doris I. The effect of micro-organisms and antibiotic substances upon viruses. | Microforms in Alexander Library. https://alexmicroforms.libraries.rutgers.edu/content/jones-doris-i-effect-micro-organisms-and-antibiotic-substances-upon-viruses.
2. The True Story of the Discovery of Streptomycin. https://www.albertschatzphd.com/?cat=articles&subcat=streptomycin&itemnum=001.
3. Jones, D. & Schatz, A. Methods of Study of Antiphage Agents Produced by Microorganisms. *J. Bacteriol.* **52**, 327–335 (1946).
4. Jones, D. The Effect of Antibiotic Substances upon Bacteriophage. *J. Bacteriol.* **50**, 341–348 (1945).
5. Schatz, A. & Jones, D. The Production of Antiphage Agents by Actinomycetes. *Bull. Torrey Bot. Club* **74**, 9–19 (1947).

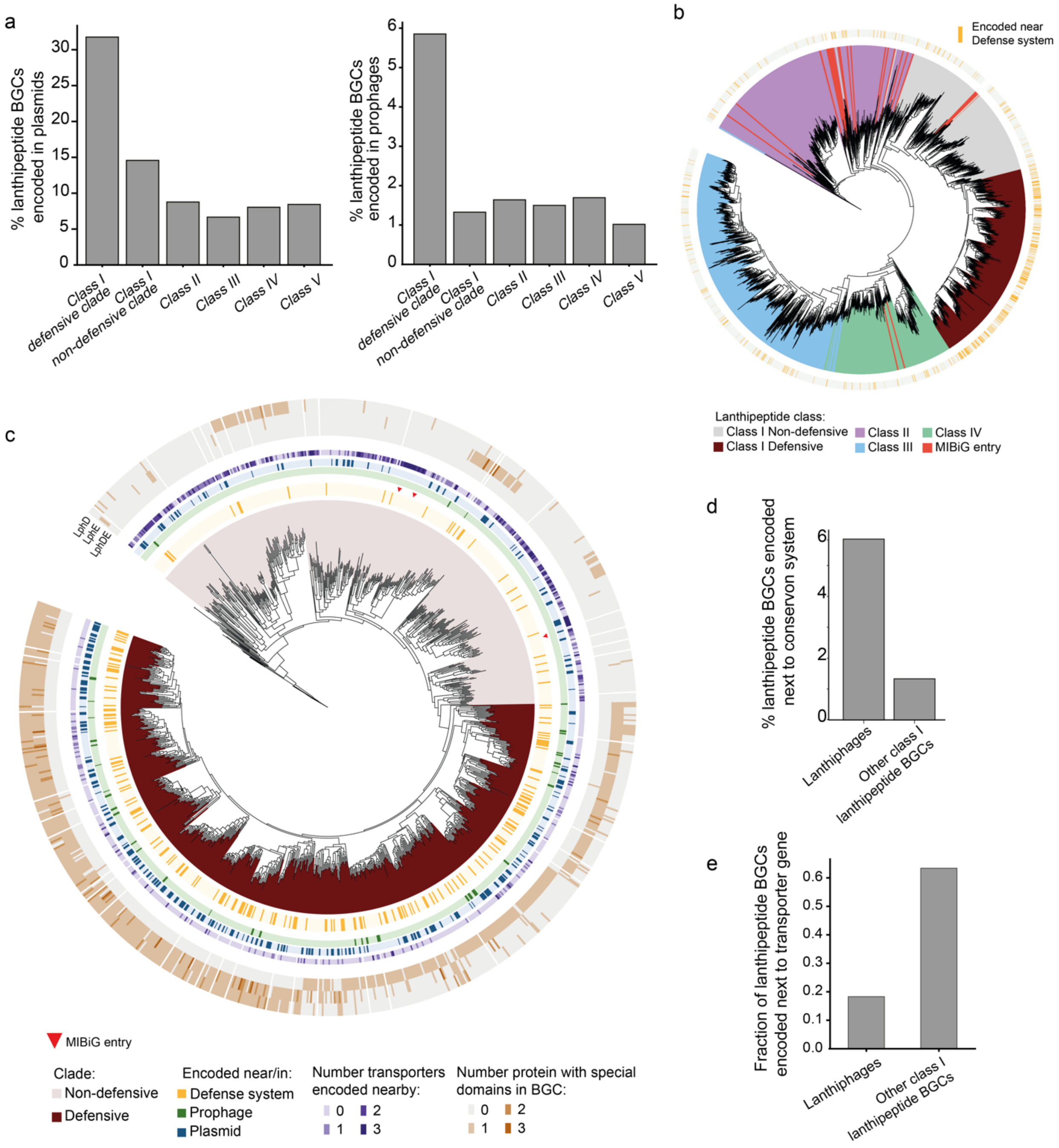


**Supplementary Figure 1** – **a**. Percentage of BGCs encoded in mobile genetic elements across different lanthipeptide classes. **b**. Phylogenetic tree of lanthipeptide BGCs using sequenced lanthionine cyclase genes encoded in actinobacterial genomes, as well as found in characterized BGCs from the MIBiG database (from diverse prokaryotic organisms). Colored ring highlights in yellow BGCs that are encoded near known defense systems. **c**. Phylogenetic tree of class I lanthipeptide BGC from Fig. 1d, with additional annotations that include the presence of accessory enzymes LphD, LphE and LphDE in the BGCs, and the number of ABC transporters encoded nearby. **d**. Percentage of lanthipeptide BGCs encoded near a conservon system. **e**. Frequency of lanthipeptide BGCs encoded near ABC transporter genes.

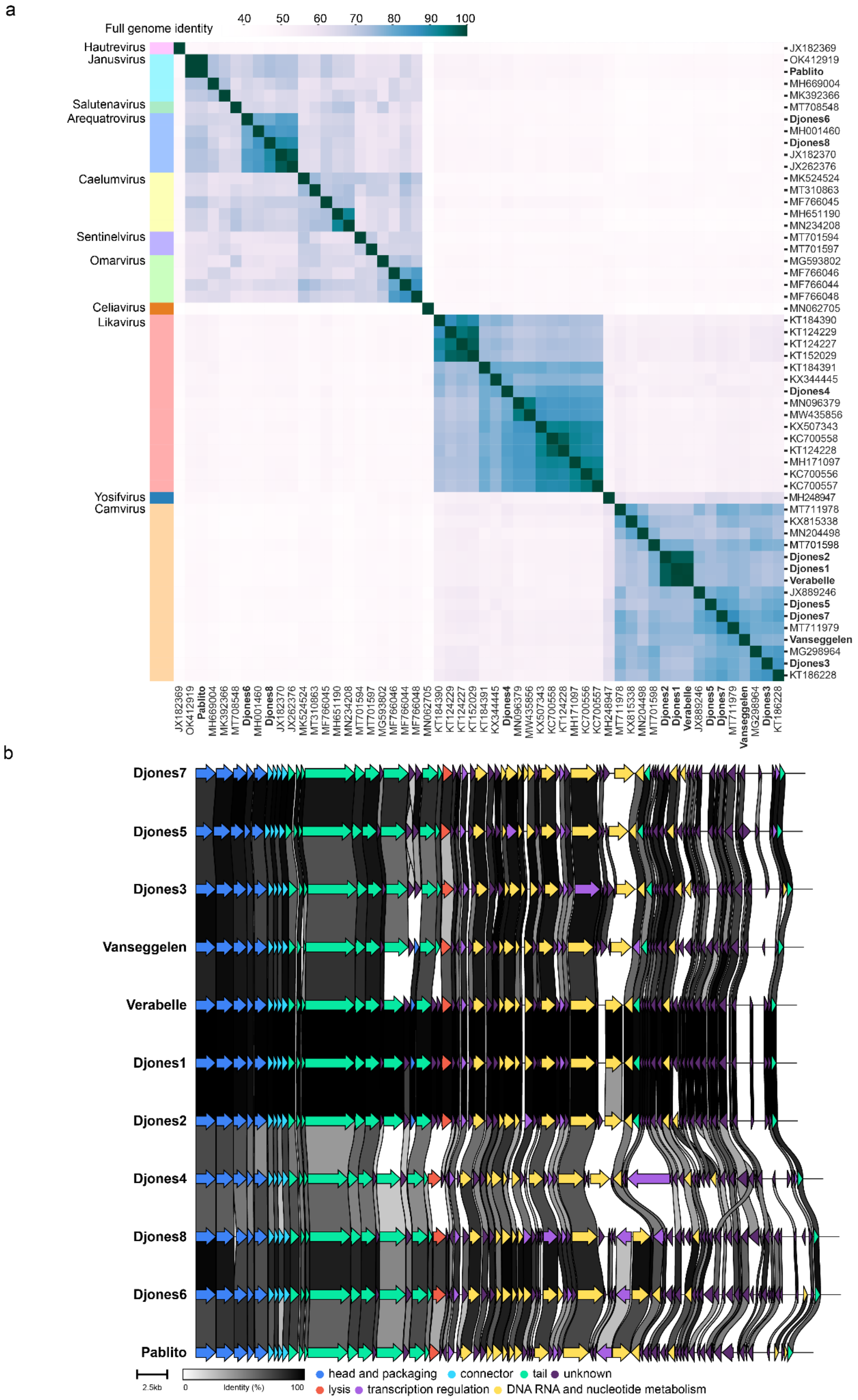


**Supplementary Figure 2** – **a**. A–Sequence identity matrix of phages from DJones collection and representative phages from the ICTV database (Arquatrovirinae). **b.** Genomic architecture of phages from the DJones collection.

a

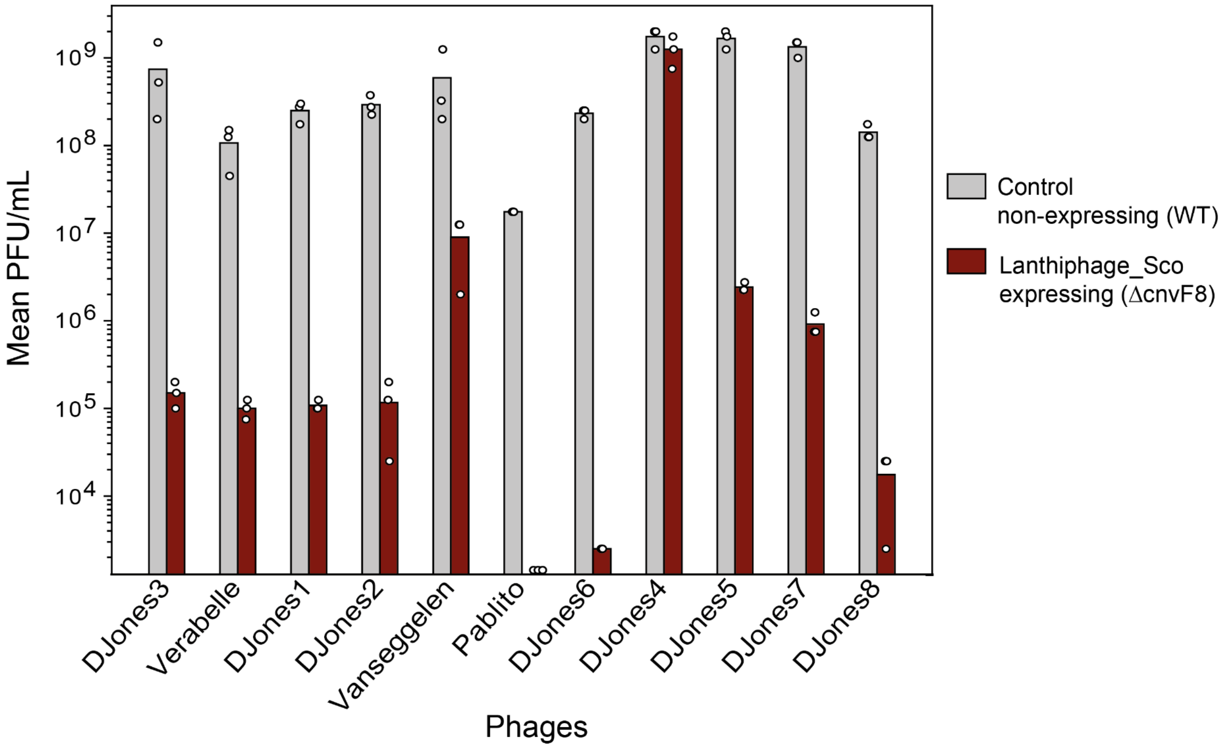


b

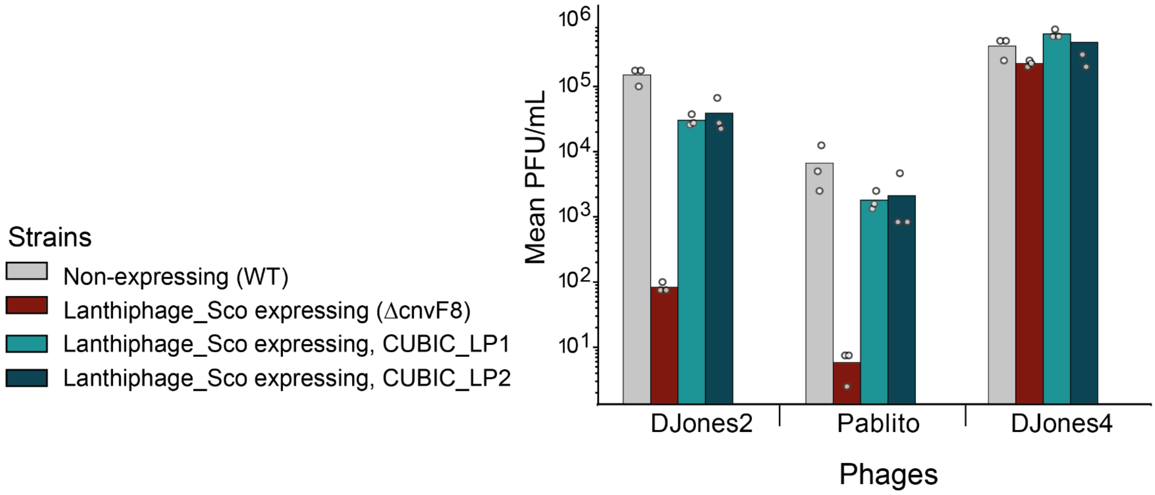


**Supplementary Figure 3** – **a.** Efficiency of plating of phages infecting *S. coelicolor* strains. Bar graphs represent an average of three biological replicates, with individual data points overlaid. **b.** Effect of knocking-down lanthiphage_Sco genes on the anti-phage resistance of the lanthiphage-expressing strain. Using the Cumate-Based Inducible CRISPRi (CUBIC) system, we designed guide RNA sequences (gRNA) to knock down the expression of both lanthipeptide genes simultaneously (strains CUBIC_LP1 and CUBIC_LP2). Bars represent an average of three biological replicates, with individual data points overlaid.

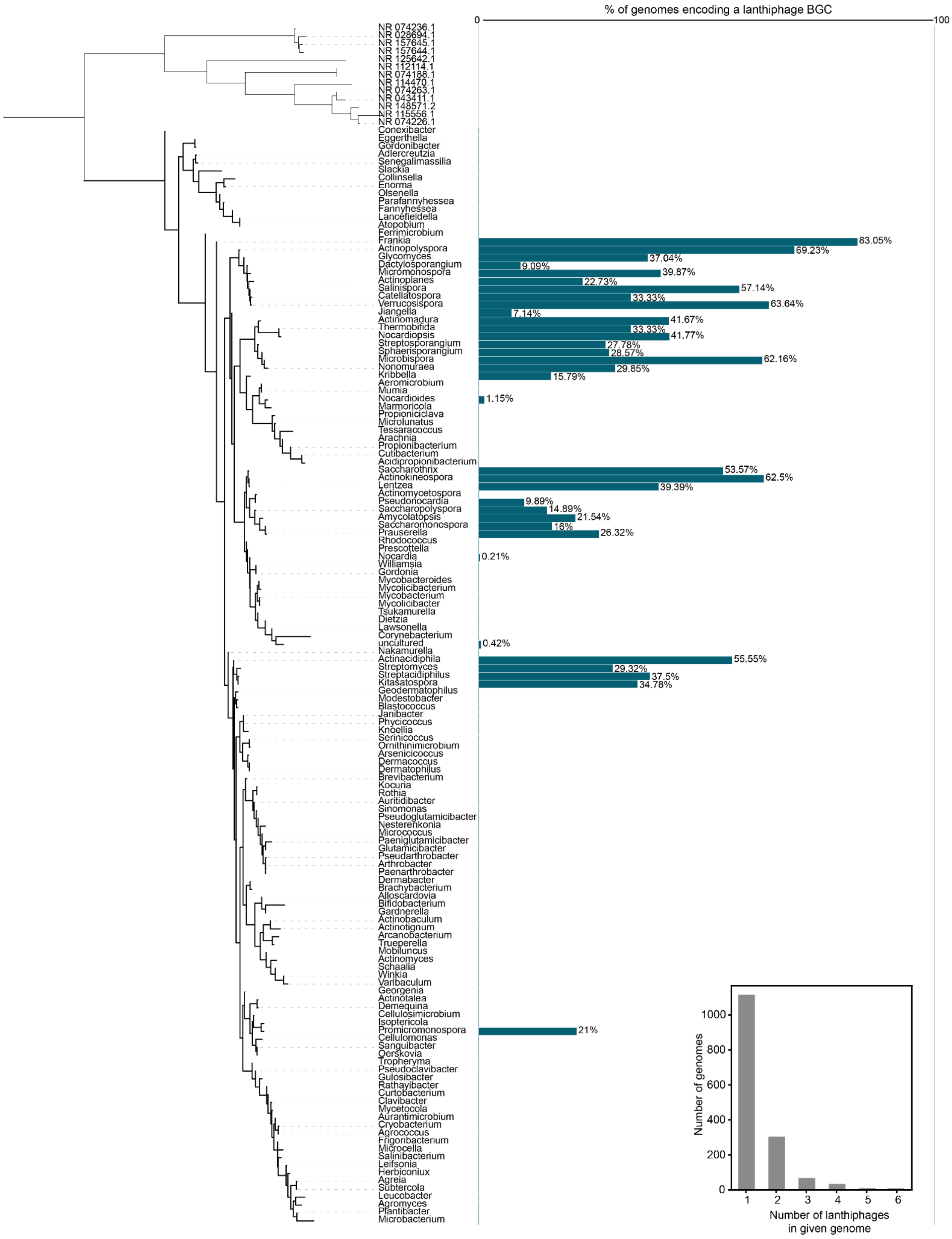


**Supplementary Figure 4** – Distribution of lanthiphages across actinobacterial genomes. Horizontal axis depicts the percentage of examined encoding lanthiphages. Numbers next to bars indicate the number of lanthiphages detected per genus. A total of 27,885 analyzed actinobacterial genomes were found without any lanthiphage. Inset: Distribution of number of lanthiphages encoded per genome.

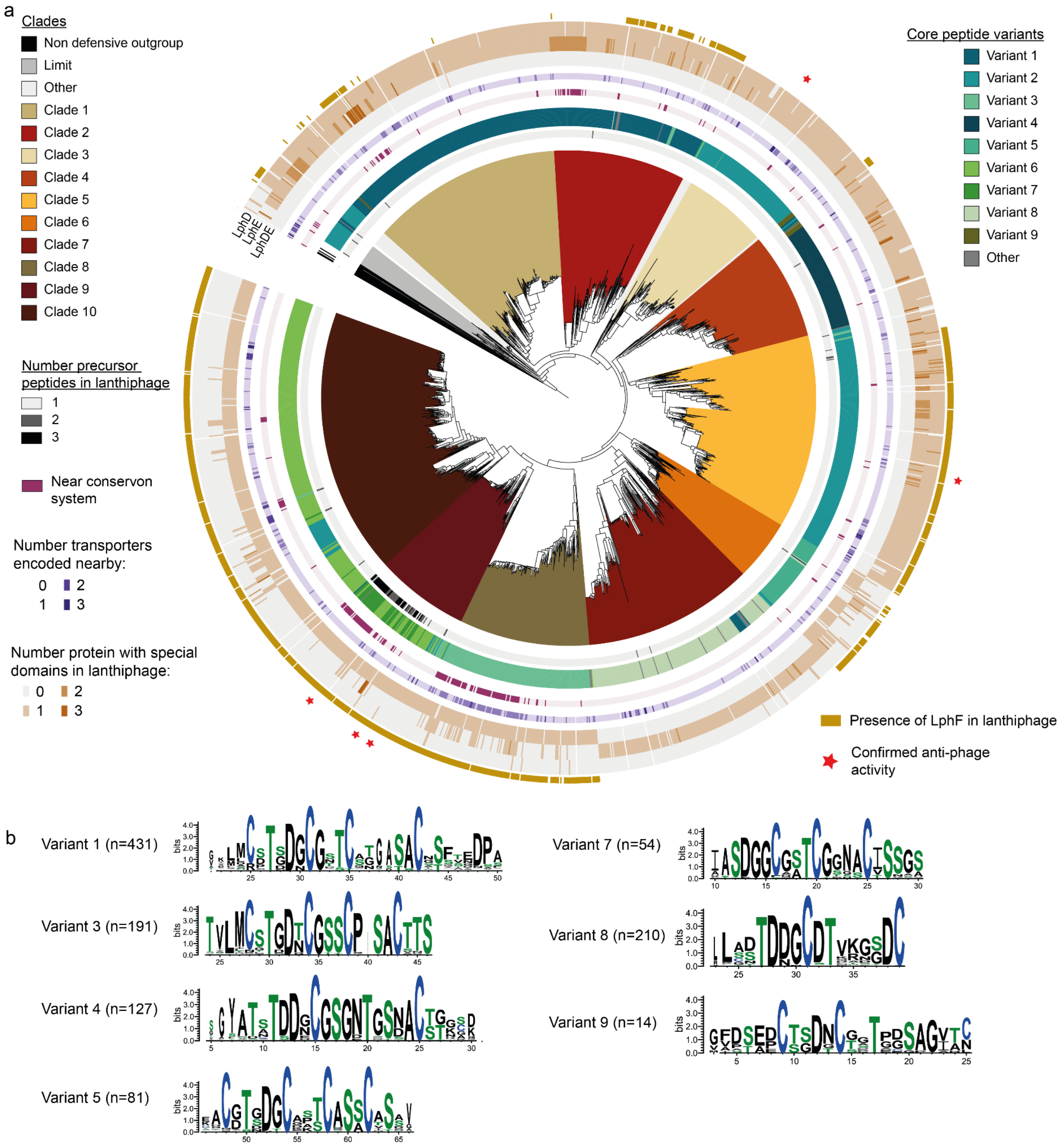


c

| Consensus formula | Variant |
|---|---|
| TXDXCXXXCXXXC | 2, 6, 8 |
| TXDXCGXXCXXXXXXC | 1 |
| TXDXCGXXCXXXXC | 3 |
| TXDXCGXGXTXXXXC | 4 |
| TDXXCXTXXXXC | 5 |
| SXXGCXXTCXXXXC | 7 |
| CXXDXCXXTXXXXG | 9 |

**Supplementary Figure 5 – a.** Phylogenetic tree of class I lanthipeptide BGCs from the lanthiphage family using their respective lanthionine cyclase sequences. Red stars indicate BGCs that were tested experimentally with confirmed anti-phage activity. **b.** Sequence logo representation of multiple sequence alignment of predicted core peptide variants, illustrating the relative frequency of conserved amino acids. Highlighted in color are residues that are post-translationally modified and participate in lanthionine ring formation: serine/threonine in green, cysteine in blue. **c.** Consensus formula describing conserved amino acids present in the different predicted core peptide variants. X, represents any amino-acid.

a

| *Name* | *Sequence* |
|---|---|
| Synthetic RBS | AAGCTTGAACAGGAGGCCCCAT |
| ErmE* promoter | TCGATCTTGACGGCTGGCGAGAGGTGCGGGGAGGATCTGACCGACGCGGTCCACACGTGG CACCGCGATGCTGTTGTGGGCACAATCGTGCCGGTTGGTAGGATCCAGCG |
| KasOp* promoter | TGTTCACATTCGAACGGTCTCTGCTTTGACAACATGCTGTGCGGTGTTGTAAAGTCGTGGCCA |

b

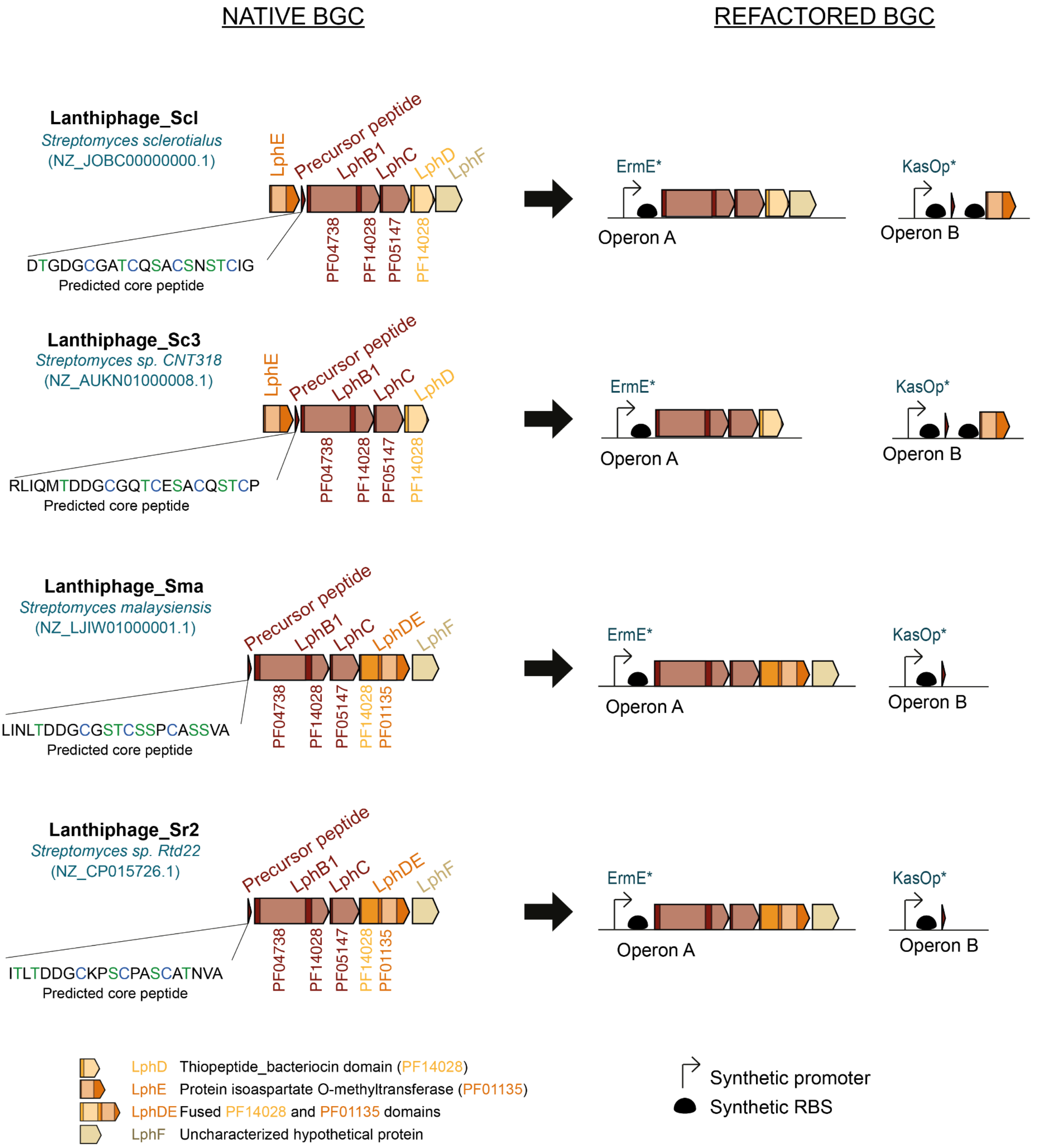


**Supplementary Figure 6** – **a.** Nucleotide sequences of synthetic promoters and RBS used in this study for refactoring lanthiphages. **b.** Schematic representation of native lanthiphages and their refactored counterparts cloned into the genome of *S. albus J1074*. Control strains contain operon A alone, while lanthiphage expressing strains contain both operons A and B.

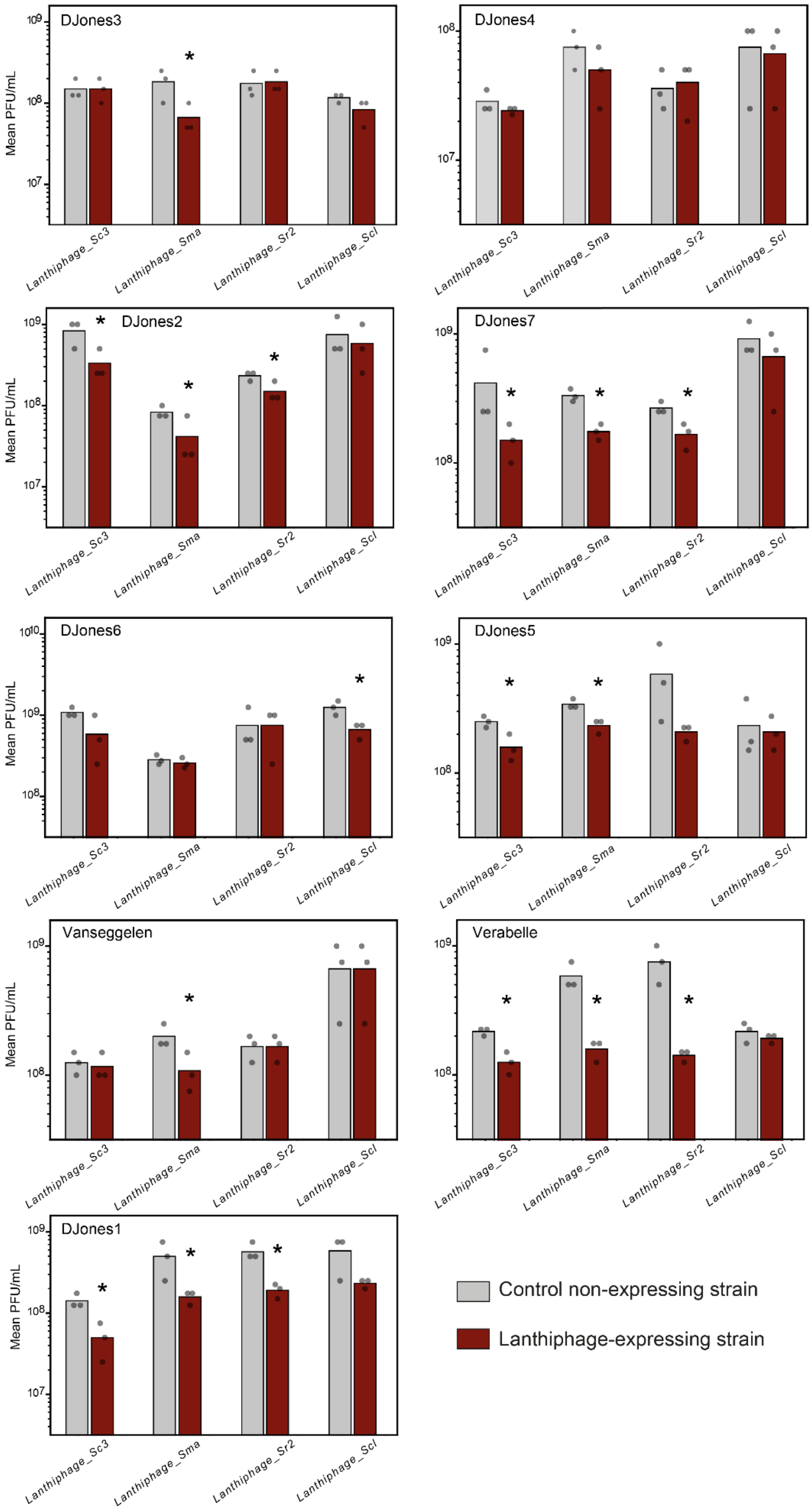


**Supplementary Figure 7** – Efficiency of plating of phages from the DJones collection on control or lanthiphage expressing strains, for all four different lanthiphages tested. Bars represent an average of three biological replicates, with individual data points overlaid. Stars indicate statistically significant reduction of PFU/mL compared to control strain (t-test, p-value<0.05).

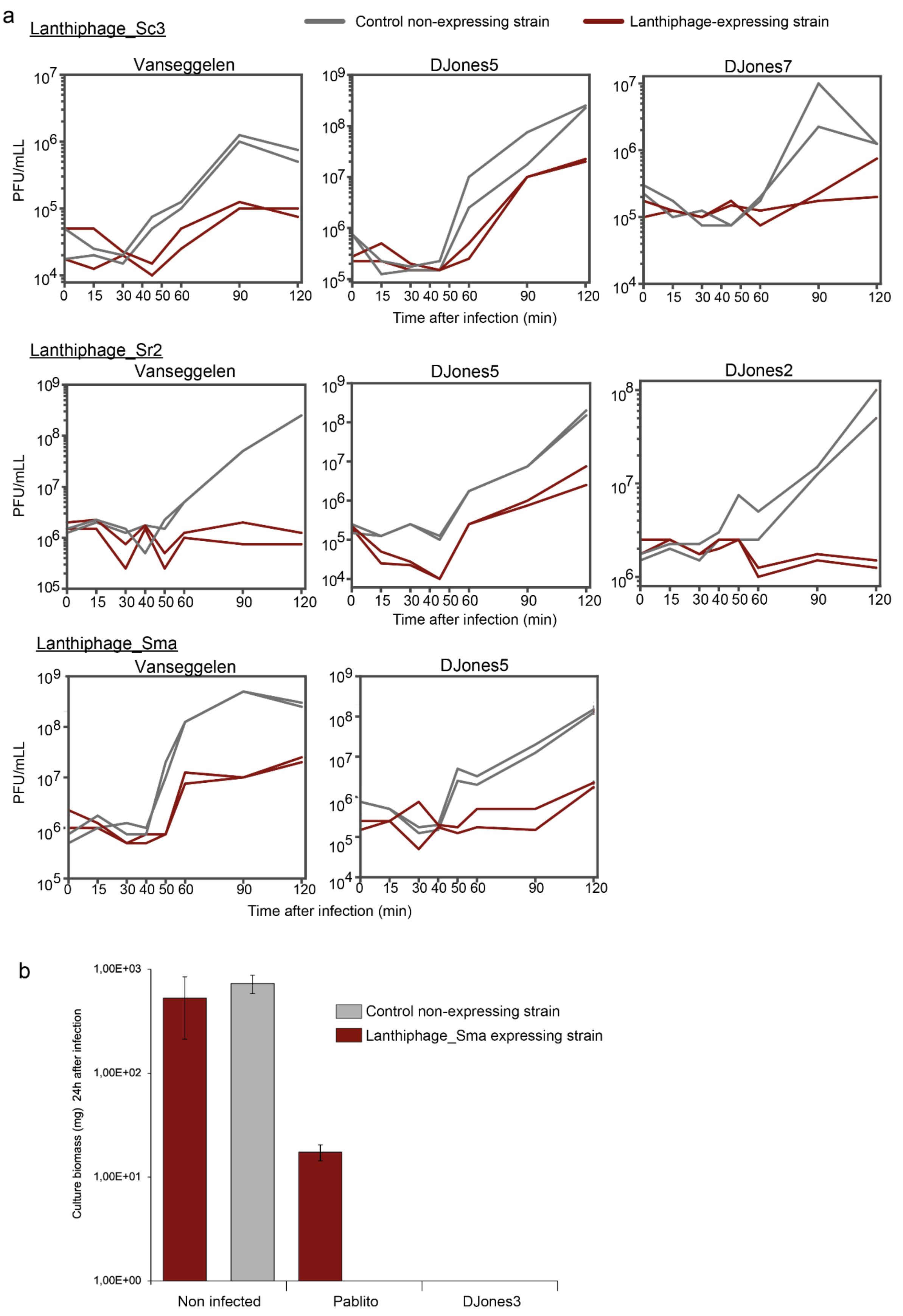


**Supplementary Figure 8** – **a.** Liquid infection assays of control and lanthiphage expressing strain infected by phages from the DJones collection at a multiplicity of infection (MOI) of 0.1. Phage plaque forming units (PFU) per mL of culture was measured over time after infection. Two biological replicates are presented as individual curves. **b.** Biomass (in mg) of cultures infected with phages at a MOI of 0.3, 24 h after infection.

a

| Medium | Strain | Mean peak area | s.d. |
|---|---|---|---|
| SFM | WT | 0 | - |
| | ΔcvnF8 | 18956.67 | 3455.41 |
| MYM | WT | 0 | - |
| | ΔcvnF8 | 35839.07 | 26440.03 |

b

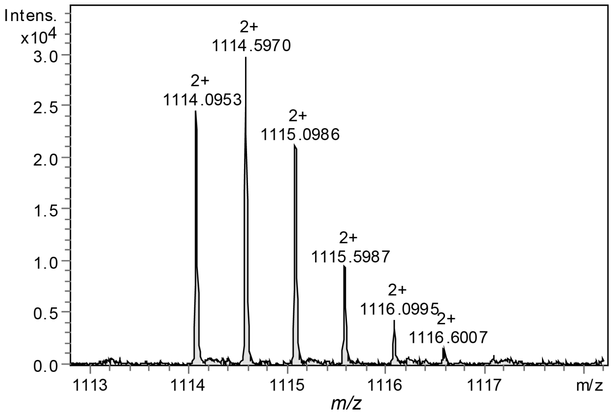


c

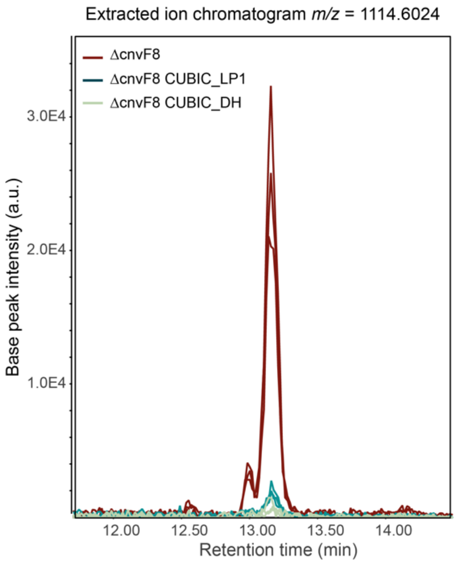


d

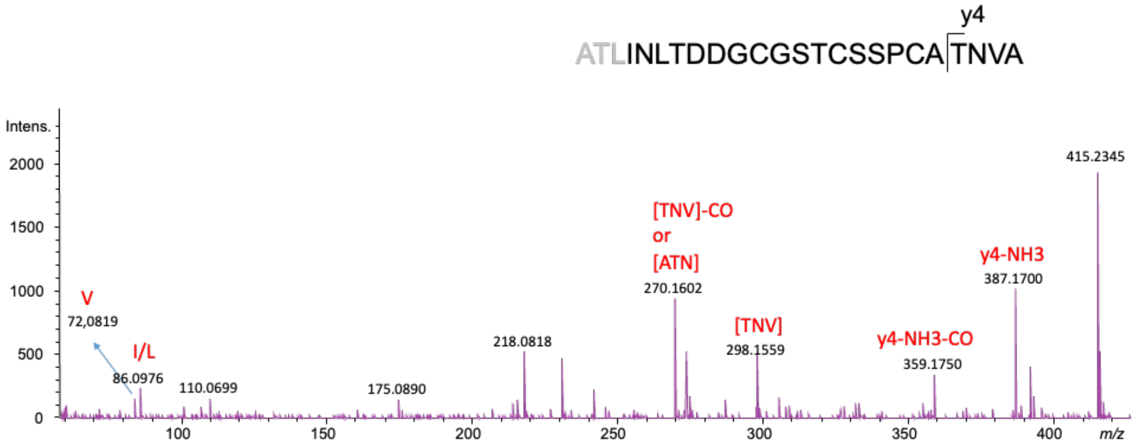


e

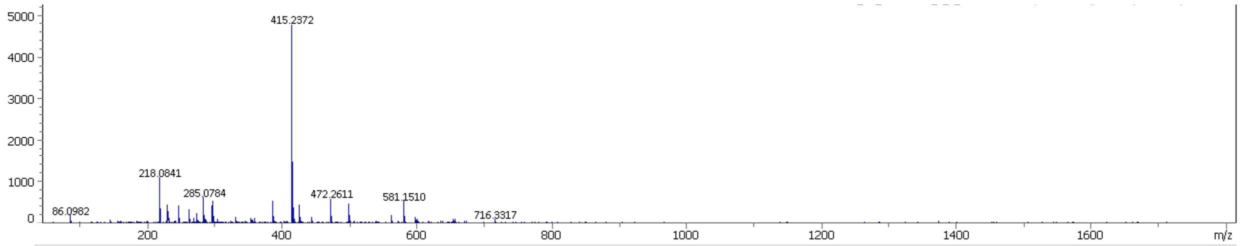


**Supplementary Figure 9** – **a.** Integrated peak area of [M+2H]$^{2+}$ ion m/z 1114.09, in cell extracts of *S. coelicolor* strains cultivated in different solid media. Mean and standard deviation (s.d.) values were calculated from three biological replicates. **b.** Mass spectrum of the [M+2H]$^{2+}$ ion m/z 1114.09. **c.** Extracted ion chromatogram of [M+2H]2+ ion at m/z 1114.09 in extracts from *S. coelicolor ΔcvnF8* strain, compared to modified strains with CUBIC_LP1 and CUBIC_DH induced with cumic acid. **d.** Tandem MS analysis of the [M+2H]$^{2+}$ ion m/z at 1114.09 corresponding to a peptide mass of 2226.17 Da. The predicted linear core peptide sequence of lanthiphage_Sco is shown. Residues in gray denote potential alternative cleavage sites. **e.** MRM fragmentation spectrum of [M+2H]$^{2+}$ ion m/z at 1114.09

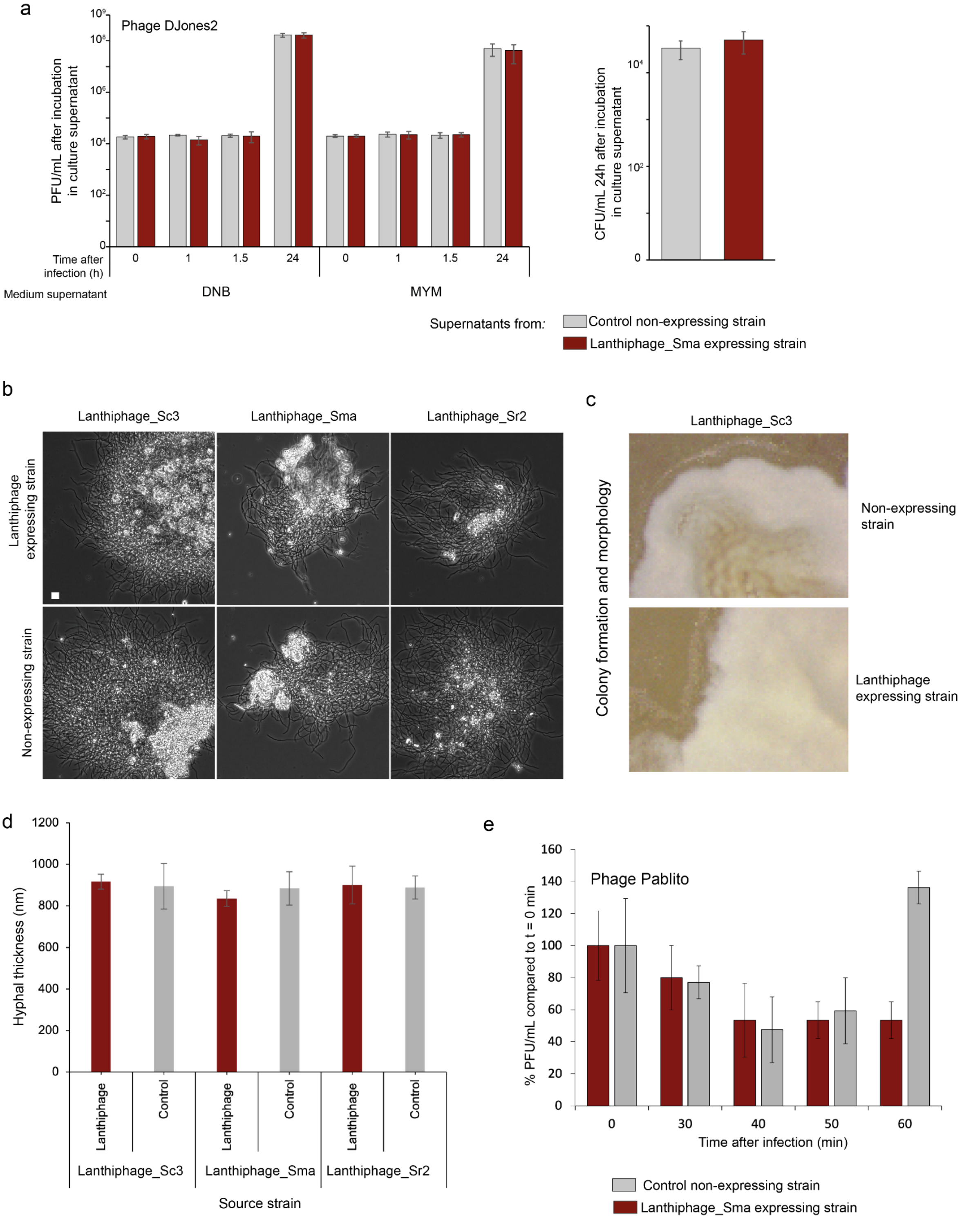


**Supplementary Figure 10 – a.** Right: Phages particles (estimated with PFU/mL) of infected WT *S. albus J1074* cultures grown in supernatants from fermentations of control and lanthiphage- expressing strains. Left: Growth of WT *S. albus J1074* cultures cultivated in supernatants from fermentations of control and lanthiphage overexpressing strains. Bar represent mean values of three biological replicates, and error bars represent standard deviation of three replicates. **b.** Microscope pictures displaying colony morphology of strains grown in liquid DNB medium, after 24 h of incubation. **c.** Colony morphology and development of control and lanthiphage expressing strains on solid MYM medium. **d.** Measured hyphal thickness of strains grown in liquid DNB medium, after 24 h of incubation. Bars represent mean values of hyphal thickness (n=6) measured with ImageJ, and error bars represent standard deviation of three replicates. **e.** Absorption assay comparing absorption of phage Pablito in control and lanthiphage-expressing strains. Bars represent mean values of three biological replicates, and error bars the standard deviation of three replicates.

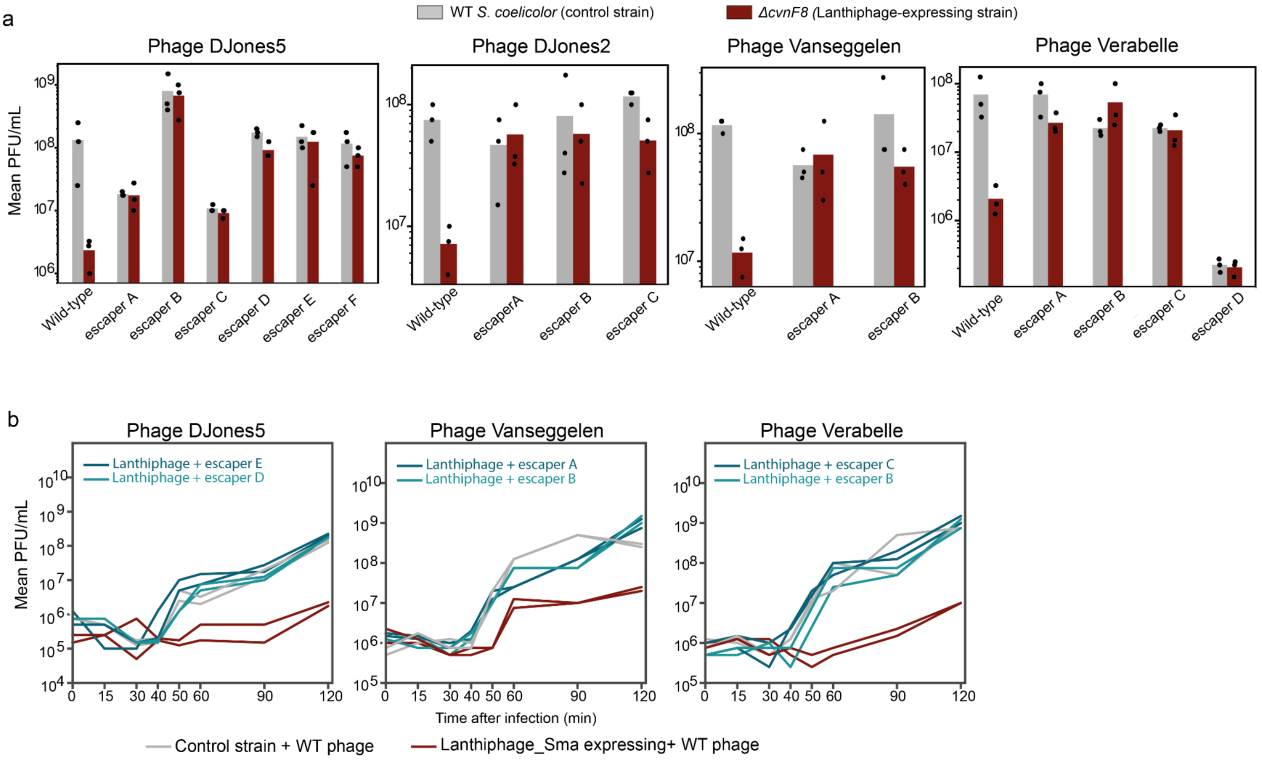


**Supplementary Figure 11** – **a.** Efficiency of plating of WT phages from the DJones collection and their respective escaper mutants, on WT or *ΔcvnF8 S. coelicolor* strains. Bars represent an average of three biological replicates, with individual data points overlaid. **b.** Liquid infection assays of control and lanthiphage-expressing strain infected by WT and escaper phages at a multiplicity of infection (MOI) of 0.1. Two biological replicates are presented as individual curves.

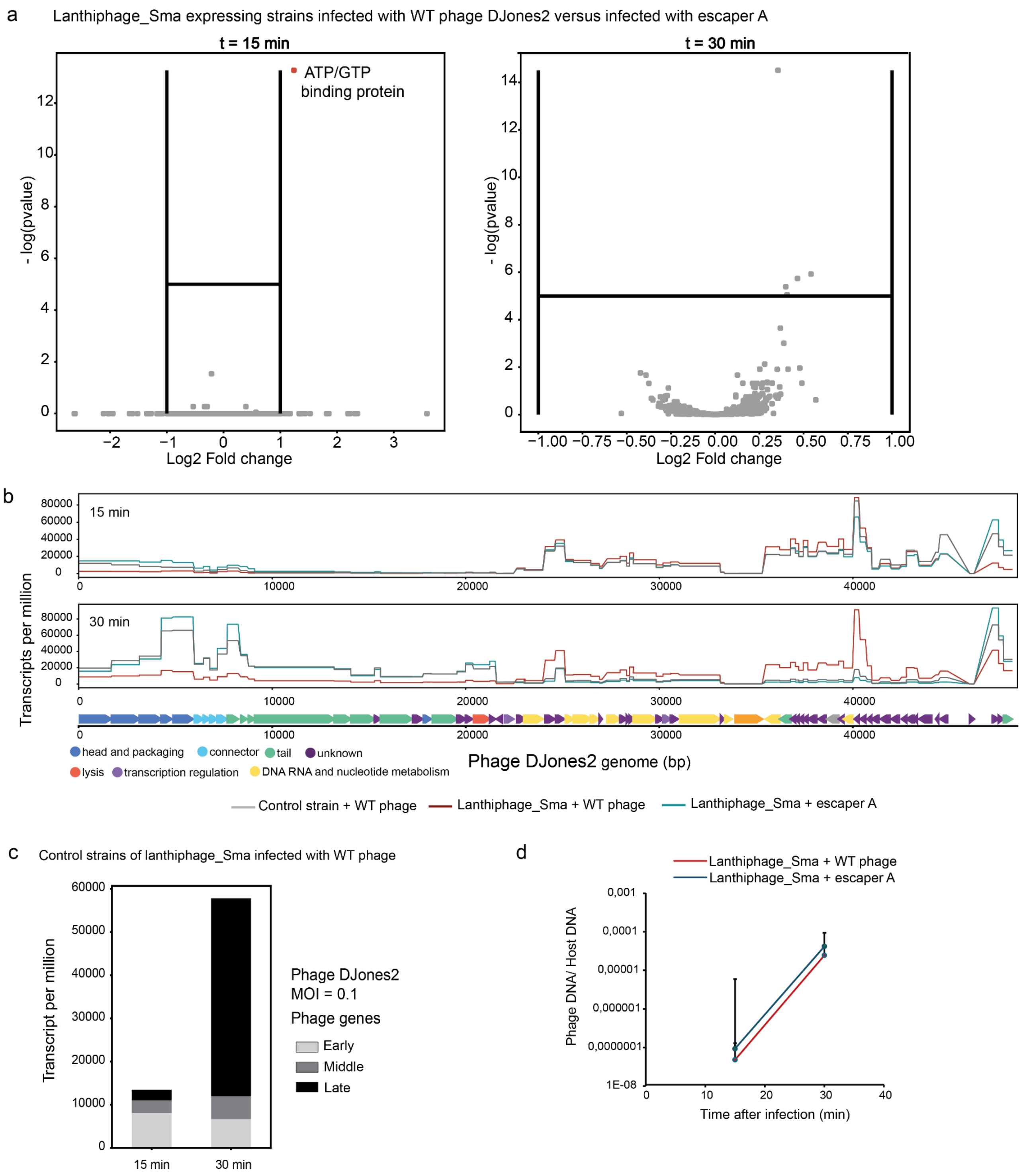


Supplementary Figure 12 – **a.** Transcripts per million of individual coding regions mapped on the genome of phage DJones2, at different time points after infection. Curves represent the average of two biological replicates per condition. **b**. Volcano plots representing all expressed transcripts of the host's genome. For every transcript, the fold change between compared conditions was plotted against the –logP values. Statistically significant differentially expressed genes, with a fold change ≥1 or ≤−1, are depicted in red and blue respectively. **c**. Level of transcription of phage genes at different time points after infection in the control strain. Bars represent the mean transcripts per million of two biological replicates, and gray stacks represent the proportion of early, middle and late phage genes transcribed. **d**. Relative concentration of phage DNA per host DNA during infection. Quantification was done by qPCR using primers amplifying a gene coding for the tail length tape measure protein (hypothetical protein) of phage DJones2 and the housekeeping gene DNA gyrase subunit B (protein id: AGI89357.1) of *S. albus*. Error bars represent standard error bars represent standard deviation of three replicates.
